# Host behavioral responses to perceived risk shape spatial disease dynamics

**DOI:** 10.64898/2026.05.25.726839

**Authors:** Dale T. Clement, Robert D. Holt, Nick W. Ruktanonchai, Omar Saucedo, Nicholas A. Kortessis

## Abstract

Host behavioral responses to disease risk are critical factors driving disease dynamics, but understanding how behavioral responses influence dynamics remains a major challenge. Coupled behavioral and epidemiological models commonly assume that hosts use population prevalence to assess disease risk. However, real-world estimates of prevalence come from data aggregated over coarse spatial scales, while transmission occurs through fine-scale contacts. Fine-scale changes in movement represent an important type of risk response because individuals must use proxies for infection risk, such as host density or environmental factors, whose relationship with actual transmission risk may vary across contexts. In this study, we examined the consequences of using different risk proxies to inform fine-scale movement and determined when imperfect proxies can cause risk-averse behaviors to increase, rather than decrease, disease transmission. We examined the effect of three risk proxies - local prevalence, local host density, and local transmission coefficient (i.e., “place”) - in the context of “simple trips”: Individuals respond to disease risk by altering travel rates from home to “away” locations and back, either leaving home less frequently (an absolute risk response) or shifting their travel to less risky away locations (a relative risk response). Absolute responses reduced prevalence more than relative responses, which were sometimes detrimental. Detrimental responses occurred when risk proxies were mismatched with the mode of disease transmission (either density- or frequency-dependent), causing individuals to either use irrelevant information or ignore pertinent information. Imperfect information thus plays a critical role in determining whether behavioral response reduces or elevates disease risk.

## 1 Introduction

Infection risk is distributed unevenly across space: local environmental conditions may inhibit or promote the growth of disease vectors (LaDeau et al. 2015; Sternberg and Thomas 2014); crowded, poorly ventilated spaces such as bars and restaurants facilitate respiratory infections (Leung 2021; Li et al. 2021; von Seidlein et al. 2021); and disease outbreaks are often spatially heterogeneous (Hagenaars et al. 2004; Lloyd and May 1996; Rodríguez and Torres-Sorando 2001). Host movement behavior — where individuals go and how long they spend there — therefore plays a critical role in mediating spatial patterns of transmission (Acevedo et al. 2015; Daversa et al. 2017; Stoddard et al. 2009). The effect of movement on transmission is a major focus of epidemiological modeling, and efforts to quantify empirically patterns of movement or space use are a critical component of understanding disease spread in both human (Arenas et al. 2020; Chang et al. 2021; Changruenngam et al. 2020; Giles et al. 2020; Stechemesser et al. 2023; Wardle et al. 2023; Zhang et al. 2022) and animal systems (Dolfi et al. 2024; Dougherty et al. 2022; Ezenwa 2004; Wilber et al. 2022). When hosts are able to perceive disease risk across the landscape and alter their movement behavior accordingly, feedback loops between disease transmission and movement behavior can result (Ezenwa et al. 2016; Hawley et al. 2021; Hoverman and Searle 2016; LeJeune et al. 2025b; Reitenbach et al. 2025; White et al. 2018).

The high salience of individual decisions to mask, social distance, and vaccinate during the COVID-19 pandemic has driven a surge of interest in linking behavioral and epidemiological modeling (reviewed in Bedson et al. 2021; Hamilton et al. 2024; LeJeune et al. 2025a; Perra 2021; Reitenbach et al. 2025). These models have examined a wide range of topics including the spread of disease awareness through the host population (Funk et al. 2009; Lobov et al. 2024; Perra et al. 2011), psychological mechanisms affecting behavior such as fear, social conformity, and exhaustion (Morsky 2025; Morsky et al. 2023; Weitz et al. 2020, reviewed in Bedson et al. 2021), and time lags in disease reporting or behavioral response (Cheng and Wu 2025; Cheng and Zou 2024; Zhang et al. 2023). Among the key insights from these studies are the observations that behavioral feedbacks can reduce peak infection and prolong an outbreak, sometimes causing multiple waves of infection (Costello et al. 2023; Morsky et al. 2023; Weitz et al. 2020; Zhang et al. 2023), and that predictive models incorporating behavior may perform better than non-behavioral models (Chen et al. 2018; Eksin et al. 2019; Osi and Ghaffarzadegan 2024; Rahmandad et al. 2022; Ryan et al. 2025).

However, many mechanistic behavioral-epidemiological models are limited in studying the effect of travel on disease spread because they focus on host decisions of whether or not to adopt a protective measure against disease, either implicitly by modeling behavioral reductions in network contacts (Arthur et al. 2021; Demirel et al. 2017; Gross et al. 2006; Machens et al. 2013; Wang et al. 2015) or transmission rate (Cheng and Wu 2025; Cheng and Zou 2024; Costello et al. 2023; Fair et al. 2024; LeJeune et al. 2024, 2025a; Mahmud et al. 2025; Rahmandad et al. 2022), or explicitly by modeling host internal beliefs or utility regarding the protective measure (Chen 2009; Eksin et al. 2017; Kumar et al. 2024; Morsky et al. 2023; Ryan et al. 2024; Schnyder et al. 2025; reviewed in Bedson et al. 2021; Funk et al. 2010; Reitenbach et al. 2025; Verelst et al. 2016; Weston et al. 2018). While such models can capture simple travel decisions — such as whether to stay home to reduce infection risk (Dashtbali and Mirzaie 2021; Fenichel 2013; Fenichel et al. 2011; Schnyder et al. 2025; Ventura et al. 2022) — or implicit changes in movement — through changes in contact networks (Eksin et al. 2017; Gross et al. 2006; Kumar et al. 2024; Maharaj and Kleczkowski 2012; Wang et al. 2015) — in general, changes in movement are extremely simplified in prior work (but see Epstein et al. 2008; Meloni et al. 2011; Nicolaides et al. 2013). Yet movement is a complex, high-dimensional behavior: individuals must make decisions about where on the landscape to move, how to get there, as well as how much time to spend before departing (Nathan et al. 2008), and this set of decisions recurs continuously amid a continuously changing risk landscape over the course of the entire outbreak (see the large literature on non-mechanistic, empirically-parameterized spatial epidemic models; Chang et al. 2021; Changruenngam et al. 2020; Koher et al. 2023; Mistry et al. 2021; Müller et al. 2021; Wilber et al. 2022). There may not be a single most protective set of decisions for reducing disease risk, and even if there is, the large space of possible decisions may lead individuals to misidentify which set of decisions is most protective (Ruktanonchai et al. 2026).

Incorporating behavior into epidemiological models requires not only choosing the behaviors that individuals adopt in response to disease risk, but also choosing the information that hosts use to assess disease risk and guide behavioral decisions. It is likely that, for both human and animal systems, hosts have incomplete or biased information about transmission risk. Many compartmental models of human disease spread that account for behavior presuppose that the information used to assess risk is some reasonably accurate estimate of disease prevalence (reviewed in Bedson et al. 2021; Funk et al. 2010; Reitenbach et al. 2025; Verelst et al. 2016; Weston et al. 2018), though imperfect information may be accounted for through time-lags (e.g., Agaba et al. 2017; Cheng and Wu 2025; LeJeune et al. 2025a; Weitz et al. 2020) or spatially-restricted information gathering (e.g., Banerjee et al. 2024; Morsky 2025; Perra et al. 2011). However, information about disease prevalence, such as case prevalence data, is often aggregated over coarse spatial scales, whereas transmission occurs through fine-scale contacts (Chang et al. 2021; Koher et al. 2023; Romeo-Aznar et al. 2022; Wilber et al. 2022). Even with coarse-level information about prevalence, local risk of infection depends on a suite of factors that individuals typically do not have information about: contact with infectious individuals, the prevalence of identifiably infectious carriers, carrier infectiousness, and persistence of the pathogen in environmental reservoirs, among others (McCallum et al. 2017; Stoddard et al. 2013; Wilber et al. 2022).

For many organisms and diseases, assessment of potential infection risk — when a person visits a restaurant, for example, or when an animal decides to forage in a particular area — requires that individuals rely on proxies for risk rather than actual risk. For respiratory diseases of humans, risk proxies may include the presence of crowds, the degree of air flow and ventilation, and municipal-level disease prevalence (e.g., Song et al. 2024). Animals may learn to associate particular environments, resources, or conspecific behaviors with pathogen risk (reviewed in Dougherty et al. 2022; Ezenwa et al. 2016; Gibson and Amoroso 2022; Stockmaier et al. 2021; Weinstein et al. 2018). Risk perception may even depend on the risk assessment of other hosts, leading to behavioral responses that are driven by the spread of disease awareness or “contagions of fear” (a long-standing topic of interest in network or agent-based disease models; Funk et al. 2009; Qiu et al. 2022; reviewed in Bedson et al. 2021; Reitenbach et al. 2025). The danger of using risk proxies to assess infection risk is that their effectiveness may be contextual: an uncrowded, but poorly-ventilated, space may pose a greater risk of COVID infection than a crowded, well-ventilated space, for example (Abdin and Mahmoud 2024; Leung 2021); or proxies that are effective for a directly-transmitted respiratory disease may be misleading for a vector borne disease (Ruktanonchai et al. 2026). As a result, risk proxies that are effective in some circumstances, may in other circumstances be weakly or even negatively correlated with actual risk. There is therefore a critical need to determine the consequences of using local risk proxies for disease transmission, and, in particular, to determine when these factors result in risk-sensitive behaviors that increase, rather than decrease, infection risk.

In this study, we examine the contextuality of movement behavior’s effect on disease transmission: how do different modes of behavioral response to disease risk interact with the mode of disease transmission and pre-disease patterns of movement to either slow down transmission of the disease or fuel its spread? We consider two key aspects of movement behavior — risk perception and risk response — that may vary from person to person or from population to population.

We use a “Simple Trip” movement model in which individuals have a fixed “home” location and move by taking “simple trips” from their home to exactly one of several discrete “away” locations before returning (see, e.g., Sattenspiel and Dietz 1995 and Citron et al. 2021, who coined the term “Simple Trip model”). This movement model is motivated by typical human movements, but can also apply to animals with a territory or home site and known foraging locations away from that site. We consider changes in movement behavior to avoid perceived infection risk that act in one of two ways: individuals either respond by spending more time at home — a site which is assumed to have negligible transmission (see below) — in proportion to average perceived risk (absolute risk response) or individuals respond by spending more time at away locations perceived as having relatively lower risk (relative risk response). Individuals may perceive risk based on any of the three components of density-dependent transmission: risk proxies associated with fixed properties of a location (i.e., local transmission coefficient; “place-based perception”), a risk proxy associated with contact rates (i.e., the density of other people at the location; “density-based perception”), and a risk proxy associated with the local disease prevalence (“prevalence-based perception”). We then measure the effect of each response and perception type on disease spread using the equilibrium global prevalence of the disease.

We consider populations that differ along along three axes of variation: 1. The overall rate at which individuals travel in the absence of disease, 2. The amount of time individuals spend at home versus away from home in the absence of disease, and 3. How strongly individuals respond to perceived disease risk. We consider infectious diseases that differ along three axes of variation: 1. The mode of disease transmission (namely, density-dependent transmission — where force of infection increases linearly with the density of infectious hosts, and frequency-dependent transmission — where force of infection is independent of host density and depends only on prevalence), 2. The strength of transmission (measured by the basic reproduction number *R*_0_), and 3. Spatial variation in transmission.

We identify which combinations of risk perception and response lead to scenarios where risk avoidance is ineffectual — or even detrimental — relative to ignoring disease risk. We find that detrimental behavioral responses tend to occur when there is a mismatch between the type of information contained in the risk proxy and the mode of transmission determining actual disease risk (either density-dependent or frequency-dependent). Detrimental responses occur both when individuals fail to use pertinent information and when individuals use irrelevant information.

## 2 Methods

We used a spatially implicit set of ordinary differential equations that allowed us to model movement activity, disease transmission, and risk perception and response. The model is summarized graphically in Figure 1, and state variables and parameters are summarized in Table 1 for ease of reference. This section describes model components according to movement, disease dynamics, risk response, and risk perception. In the main text, we present a simplified three-patch model. A more general *n* patch formulation of the model is given in Supplement 1.

**Table 1:**
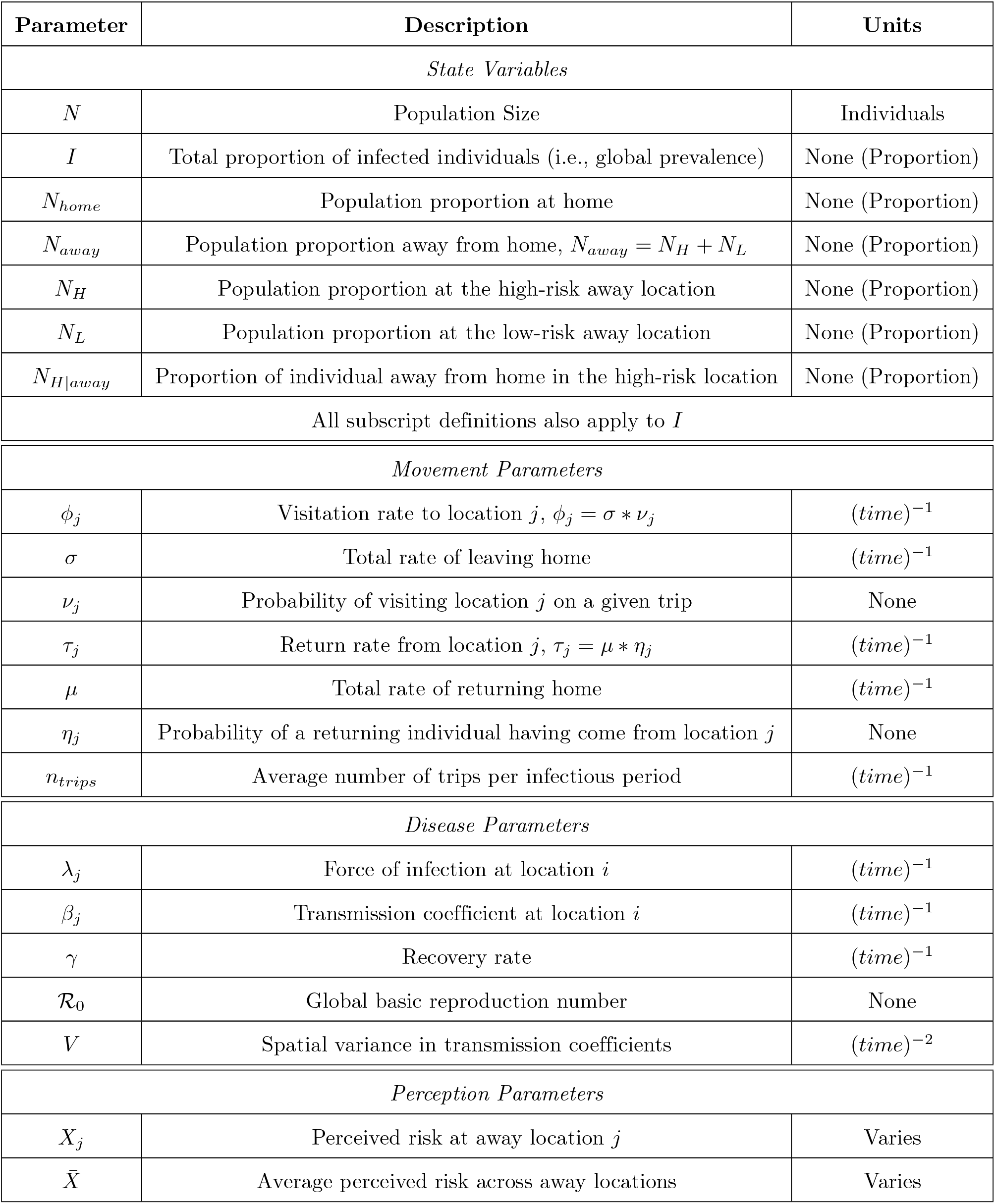
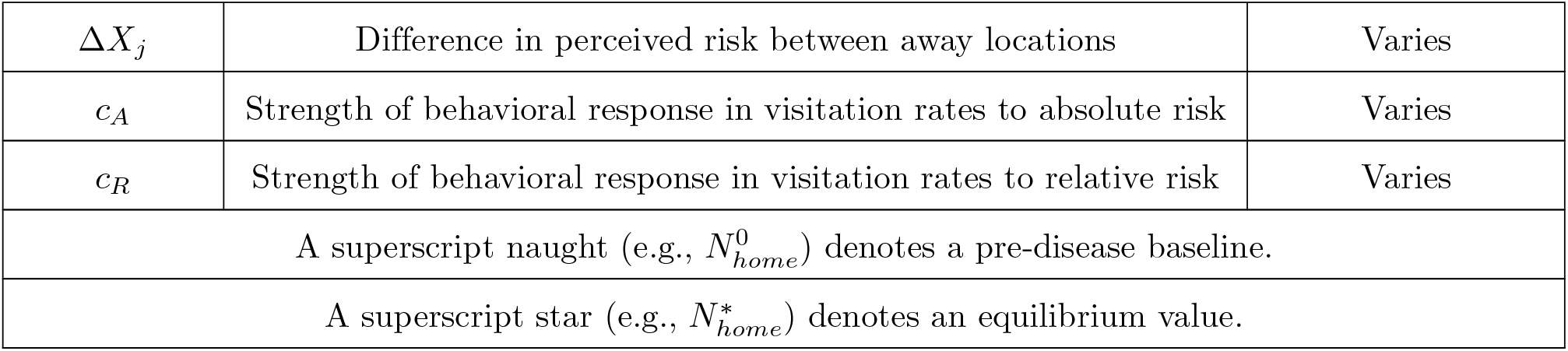
A description of parameter notation used in the model. Note that the products 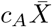 and *c*_*R*_Δ*X*_*j*_ are always unit-less and the units of *c*_*A*_ and *c*_*R*_ therefore depend on the perception variable

**Figure 1:**
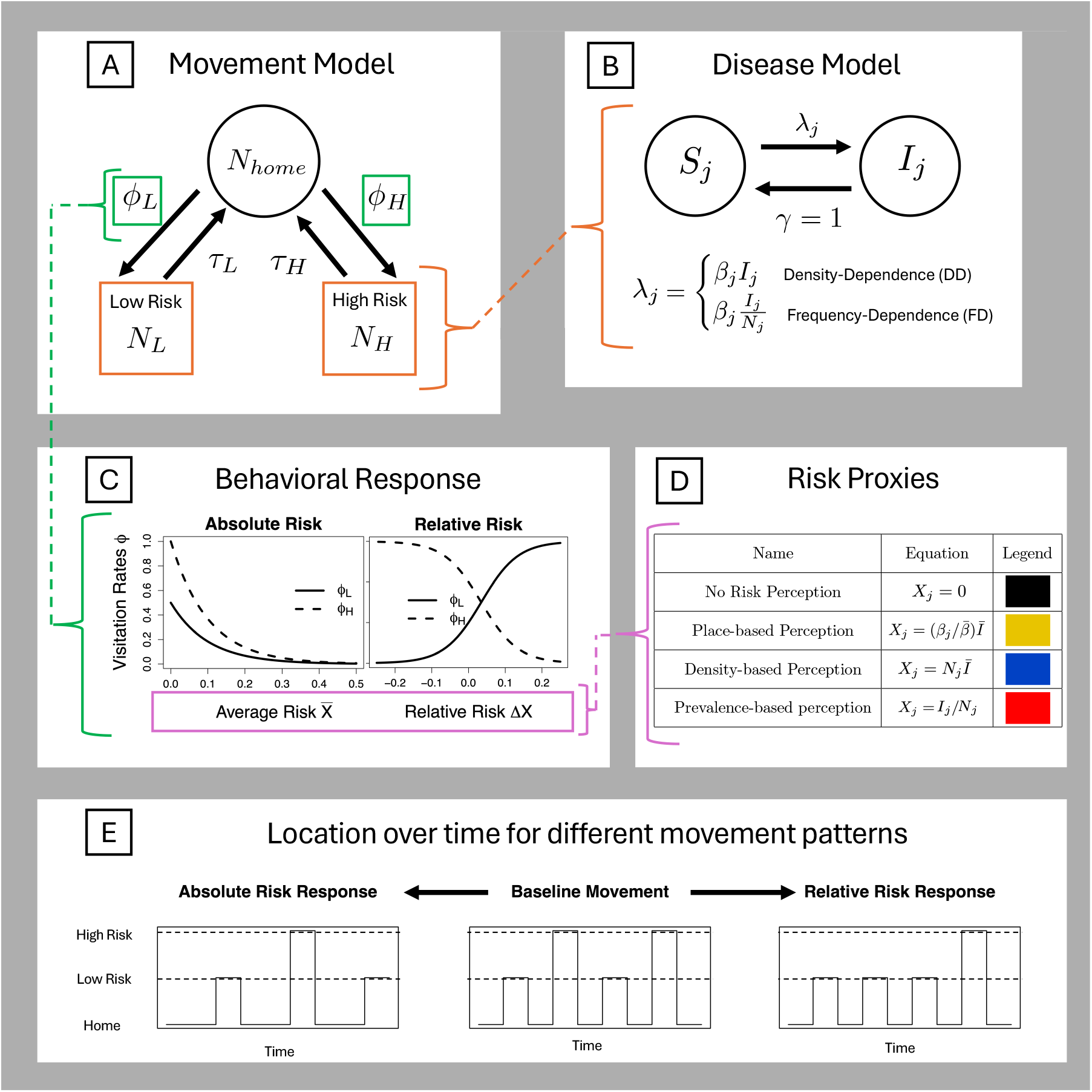
A schematic of the model structure. A) Movement follows a Simple Trip model with one home location (no disease transmission) and two away locations (one with high transmission coefficient, the other with low transmission coefficient). B) Within each location, the disease follows a Susceptible-Infectious-Susceptible (SIS) model and we consider both density-dependent and frequency-dependent transmission. C) The rate at which individuals leave home decreases with average perceived risk, while the probability of visiting each away location changes with the perceived risk differential between the two locations. D) Individuals use one of three proxies (or no proxy) to assess disease risk in each away location: local transmission coefficient (i.e., “place”), local density, local prevalence. Each proxy has an associated color used to represent it in the results figures. E) A conceptual diagram of how movement patterns change due to absolute risk response and relative risk response.

### Disease-free movement

Consider a sufficiently large population of *N* hosts divided among three discrete locations: a home location with *n*_*home*_ individuals, a low-risk away location with *n*_*L*_ individuals, and a high-risk away location with *n*_*H*_ individuals. We may, without loss of generality, normalize by the total population size so that *N*_*home*_ = *n*_*home*_*/N* denotes the proportion of individuals at home, *N*_*L*_ = *n*_*L*_*/N* denotes the proportion of individuals in the low-risk location, and *N*_*H*_ = *n*_*H*_*/N* denotes the proportion of individuals in the high-risk location.

Because we were interested in modeling the response of frequent, local movement to disease risk, the “home” location, rather than representing a single physical location, is best interpreted as the state of being in one’s own domicile. The two away locations then represent any two points of interest that an individual may visit in their day-to-day travels: shops, restaurants, grocery stores, parks, work places, etc. The away locations are distinguished by the degree to which they facilitate the spread of disease (in our “canonical” example, a respiratory infection). This could be differences in ventilation (e.g., between parks and restaurants), hygiene policies (e.g., in masking policies, or other behavioral shifts), or any other intrinsic feature that affects disease transmission.

In the absence of disease (denoted with a superscript 0), individuals at home visit away location *j* ∈ {*H, L*} at a rate 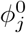 . Visitation rates are composed of the total rate at which trips are taken, *σ*^0^, times the probability of visiting location *j* given that a trip occurs, 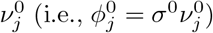 . Individuals return from location *j* at a rate 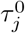, which may similarly be expressed as the total rate *µ*^0^ at which residents return from trips, irrespective of origin, times the probability 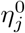 that a returning individual came from location *j*, (i.e., 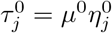). This yields the Simple Trip movement model

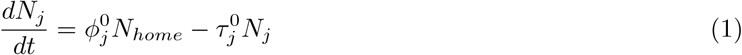

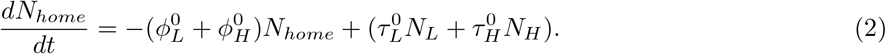

The disease-free equilibrium densities of the Simple Trip movement model are 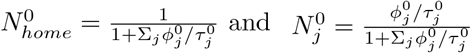 .Because 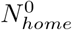 and 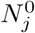 are proportions of individuals, they can also be interpreted as the average fraction of time individuals spend at home and in each away location (Sattenspiel and Dietz 1995). We show in Supplement 2 that the average number of trips per unit time is 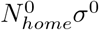, or the average fraction of time spent at home 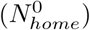divided by the average time spent at home between trips (1*/σ*^0^).

### Disease Model

We consider a susceptible-infected-susceptible (SIS) compartmental disease model coupled to the Simple Trip movement model. Because the base movement model is written in terms of *N*_*j*_, we track changes in *N*_*j*_ instead of *S*_*j*_ (with *N*_*j*_ = *I*_*j*_ + *S*_*j*_). Our Simple Trip SIS model assumes that contact rates at home between infected and susceptible individuals are sufficiently low compared to those away such that the transmission rate at home can be assumed zero (*β*_*home*_ = 0). We relax this assumption in Supplement 5 and discuss its implications in the Discussion. Under this assumption, the model is written as

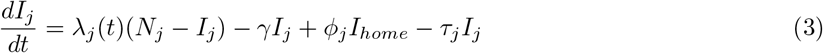

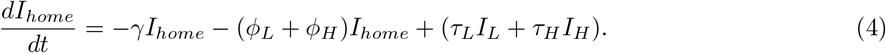

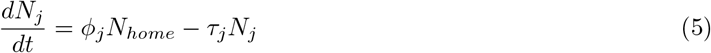

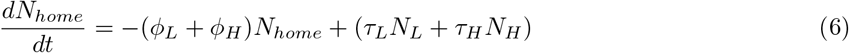

where *λ*_*j*_(*t*) is the force of infection in away location *j* at time *t*, and *γ* is the rate at which infectious individuals become susceptible. We consider both density-dependent (*λ*_*j*_(*t*) = *β*_*j*_ *I*_*j*_) and frequency-dependent 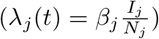 forms for the force of infection, where *β*_*j*_ is the location-specific transmission rate. Without loss of generality, we set *γ* = 1, which is equivalent to scaling time so that all rates are given in units per infectious duration (1*/γ*), and all transmission coefficients *β*_*j*_ are given in units per infectious duration.

### Risk Response

We model behavioral responses to disease as changes in the visitation rates *ϕ*_*j*_ to away locations *j* ∈ {*H, L*}. The generalized model presented in Supplement 1 allows behavioral responses to include both reductions in visitations rates (less frequent trips) *ϕ*_*j*_ and increases in return rates *τ*_*j*_ (shorter trips). The effects of risk avoidance on global prevalence (*I*^*^) were similar regardless of whether individuals reduced visitation rates or increased return rates. We therefore present only risk responses that reduce visitation rates to high risk areas.

Baseline movement rates, 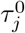 and 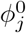, may be interpreted as the average preference for different locations in the absence of disease, and deviations from the baseline preference represent individual avoidance based on (perceived) disease risk. Because home locations as assumed to have zero transmission, staying home always represents the most risk-averse behavior.

Individuals adjust their visitation rates based on perceived risk *X*_*j*_, which represents the information individuals use to assess disease risk in away location *j*. Just as baseline visitation rates may be separated into total visitation rate and visitation probabilities 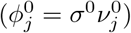, we construct the risk response function *ϕ*_*j*_(*X*_*j*_) so that it is separable into two components: changes in total visitation rate as a function of average perceived risk 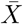 across away locations, which we define as “absolute risk avoidance” 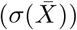, and change in visitation probabilities as a function of perceived risk at location *j* relative to the average risk, 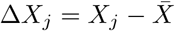, which we define as “relative risk avoidance” (*v*_*j*_(Δ*X*_*j*_)). We model visitation rate as an exponential function of *X*_*j*_ so that total risk avoidance (i.e., the dependence of *ϕ*_*j*_ on *X*_*j*_) may be written as the product of absolute and relative risk avoidance: 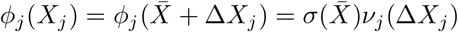 .

We may then write visitation rate as

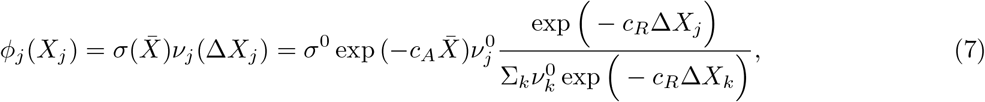

for *k* ∈ {*H, L*}, where *c*_*A*_ controls the absolute decrease in visitation rate with perceived disease risk and *c*_*R*_ controls the relative decrease in visitation rate with perceived disease risk. The change in *v*_*j*_ with *X*_*j*_ is standardized so that *v*_*H*_ + *v*_*L*_ = 1.

If individuals spend different amounts of time in each location, behaviorally-mediated changes in relative visitation rates (*v*_*j*_) have the potential to influence the total time spent away, thereby mimicking the effect of absolute behavioral response by modifying the time spent at home. To ensure that the total time spent at home remains constant while considering only relative risk response (i.e., *c*_*A*_ = 0 and *c*_*R*_ *>* 0), we adjusted the return rates as follows:

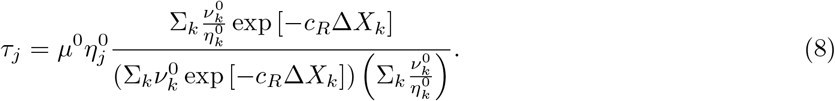

This ensures that the total fraction of time spent at home is constant for all *c*_*R*_.

To verify that this is the case, observe that the equilibrium fraction of time spent at home, for constant *X*_*j*_, is

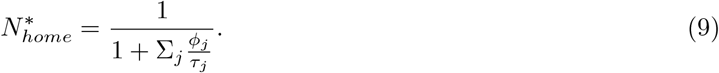

From equations 7 and 8, the ratio of visitation to return rates, for constant *X*_*j*_, is

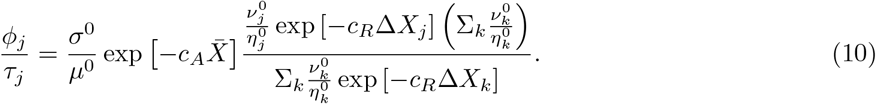

Using equation 10 in equation 9 yields

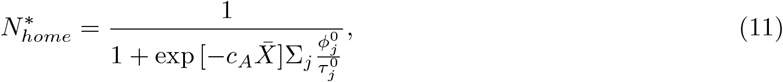

which does not depend on the relative scaling parameter *c*_*R*_. Note also that the equilibrium fraction of time spent away from home is then

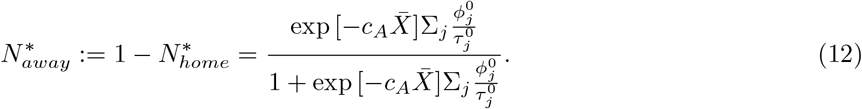

As such, the fraction of time spent in away location *j* conditional on being away is

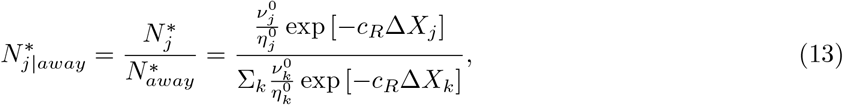

demonstrating that relative risk response (*c*_*R*_ *>* 0) shifts only the proportion of time individuals spend in away locations of their total allotment of time away from home.

As noted above, the scaling parameters *c*_*A*_ and *c*_*R*_ control the strength of each type of behavioral response. Setting one or more of these parameters to zero has the effect of “turning off” a given response type. This enables us to study each response type in isolation by setting the parameters controlling the other responses to zero. We assume that the scaling parameters are positive, such that greater perceived risk results in individuals spending less time in locations perceived to be risky.

### Risk Perception

We construct each risk proxy as one of three underlying types of information that combine to form the density-dependent transmission rate *β*_*j*_*I*_*j*_. These sources of information are disease prevalence 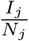, host density (i.e., the proportion of individuals *N*_*j*_), and local transmission coefficients *β*_*j*_. Because disease-dependent behavior requires perceived risk to be a function of infections, we assume that each risk proxy also incorporates information about global disease prevalence *I*, which implies that individuals, regardless of the local risk proxy they respond to, nonetheless have some sense of the overall burden of disease in the population.

The three core risk proxies are thus prevalence-based risk perception 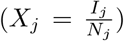, density-based risk perception 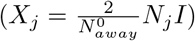, and place-based risk perception 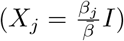. We scale density-based perception by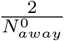 to ensure that perception is consistent across different spatial structures and baseline movement rates. Without scaling *X*_*j*_ by the number of away locations, in this case *K*_*away*_ = 2, our definition of *N*_*j*_ as a proportion would cause perceived density-based risk to decrease with the number of away locations in the general Simple Trip formulation (see Supplement 1). Further, without scaling by 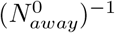 populations that spend more time at home in the absence of disease would perceive less risk when a disease arises.

It should be noted while some positive relationship between global prevalence and risk is necessary to ensure risk responses are not exhibited prior to the introduction of the disease, this relationship need not be linear. One could write a more generalized form of, for example, place-based risk-perception as *X*_*j*_ = *β*_*j*_*f* (*I*), where setting this function as an indicator variable (e.g., *f* (*I*) = **1**_*I>a*_) would then cause individuals to exhibit a binary response triggered once disease risk reaches a minimum threshold (a commonly employed functional form; reviewed in Verelst et al. 2016).

There are any number of reasons why an individual might rely on one type or source of information to the exclusion of others. For example, individuals may not consider *β*_*j*_ if individuals do not distinguish risk among indoor locations (e.g., do not pay particular attention to type or degree of ventilation) and only consider how crowded each location is. As mentioned in the introduction, local prevalence information may not always be available and, early during the outbreak of a novel disease, information about local transmission may not be available, either. More generally, however, the goal of partitioning risk into underlying types of information and examining them separately is to develop a theoretical understanding of how different kinds of partial information affect individual behavior.

### Numerical Analysis

We numerically analyzed our three patch model by considering two response modes (setting each of *c*_*A*_ and *c*_*R*_ positive while holding the other at zero), three information types (prevalence-based, density-based, and place-based perception), and two modes of disease transmission (frequency-dependent and density-dependent) leading to 12 combinations of response mode, information type, and transmission mode. We manipulated five key parameters to determine their effect on *I*^*^ for each of the 12 combinations. First, we varied total movement rate, measured as number of trips per infectious duration *n*_*trips*_, on the log scale from 10^−3^ to 103. Movement rate plays a critical role in the epidemiological interpretation of the model, with faster movement rates best describing short-term, local travel and slower movement rates best describing less frequent, longer-distance travel. The range from 10^−3^ to 10^3^ represents a gradation between extremes of very fast and very slow movement, which are limiting cases where timescale separation allows for some degree of analytic insight (see Supplement 4).

Second, we varied the pre-disease proportion of time spent in the low transmission location 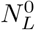 by varying the probability of visiting the low-risk location 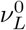 from 0.01 to 0.99. Pre-disease behavior is important because it serves at the baseline against which the effectiveness of a risk response is determined. Equations converting movement rate, *n*_*trips*_, and pre-disease proportion of time spent at home, 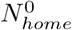, to total visitation and return rates, *σ*^0^ and *µ*^0^, are given in the Supplement 2.

Third, we varied the global basic reproduction number *R*_0_ — defined as the population-weighted average of local values of *R*_0_ when the population has settled into its disease-free behavioral equilibrium — from 1 to 8, ranging from a disease that barely persists in the population to a highly infectious disease. Fourth, we varied spatial variation in transmission coefficients — defined as the standard deviation 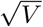 of the transmission coefficients *β*_*L*_ and *β*_*H*_ — from 0 to 2*R*_0_. Equations converting values of global *R*_0_ and transmission standard deviation 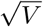 to values of *β*_*L*_ and *β*_*H*_ are given in Supplement 3. Finally, we varied the strength of behavioral response *c* from 0 to 20, ranging from insensitive to highly sensitive responses to risk.

Because we were interested in modeling responses in frequent, local movement to respiratory disease risk, we chose as baseline parameters relatively fast movement (*n*_*trips*_ = 10), individuals who spend most of their time away from home 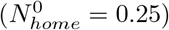, moderate transmissibility (*R*_0_ = 2), and moderate spatial variation in transmission 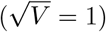. Common respiratory pathogens have *R*_0_ in the range of 2–3 and infectious periods in the range of 3–10 days (see table 3 in DeWitt et al. 2024) such that 10 trips per infectious period corresponds to 1–3 trips per day. In the absence of disease, individuals visit the high risk location twice as frequently as the low risk location (*v*_*L*_ = 1*/*3, *v*_*H*_ = 2*/*3), emphasizing the tradeoff between avoiding disease risk and fulfilling baseline preferences. We ensure individuals spend similar times in each location on any given trip by setting relative return rates as *η*_*H*_ = *η*_*L*_ = 1*/*2. Individuals exhibit behavioral responses to disease risk of intermediate strength (*c* = 10). To assess the sensitivity of our results to movement speed, we also run the above analyses for *n*_*trips*_ = 0.1 (see Figure S4), but note that the qualitative conclusions are little changed at this slower movement rate.

## 3 Results

### How does disease prevalence depend on risk response?

Across all the cases we considered, spending more time at home (i.e., absolute risk avoidance) was consistently more effective at lowering global prevalence than moving from more to less risky away locations (i.e., relative risk avoidance). The benefits of staying home (as compared to shifting to safer away locations) increased with more transmissible diseases (i.e., larger *R*_0_; Figure 2a,b for density-dependent transmission; Figure S1a,b for frequency-dependent transmission).

**Figure 2:**
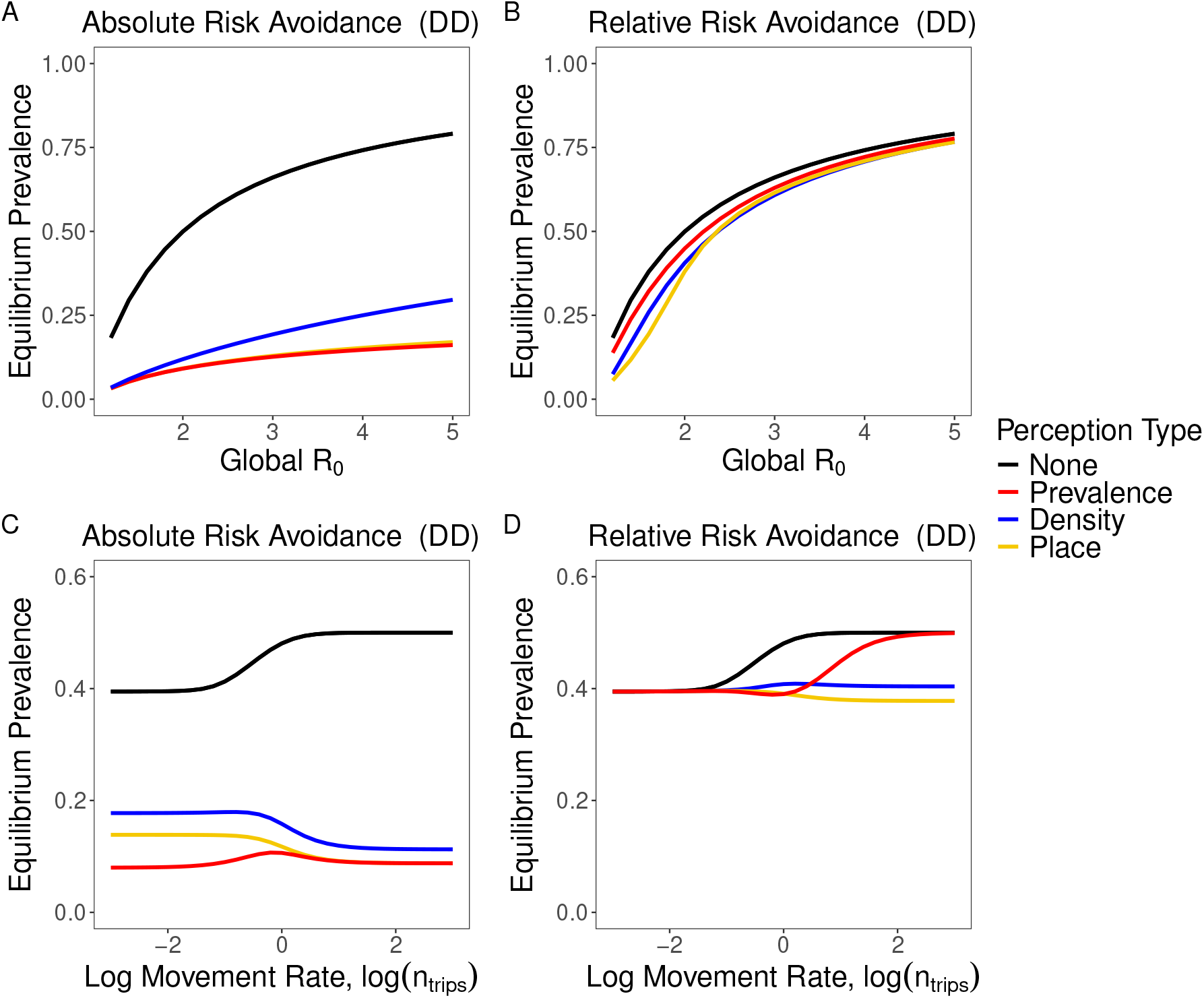
Equilibrium disease prevalence when risk-induced changes in travel behavior do not occur (black), rely on information about local prevalence (red), rely on information about local host density (blue), or rely on information about the transmission coefficients *β* in each location (gold). Top row: effect on equilibrium prevalence due to changes in *R*_0_, holding spatial variance in transmission constant. Bottom row: effect on equilibrium prevalence due to changes in movement rate (log number of trips per infectious duration). Left column: individuals respond to disease risk by increasing the proportion of time they spend at home, 1*/σ* (absolute risk response). Right column: individuals respond to disease risk by increasing the proportion of time they spend in the less risky away location, with time at home held constant. Transmission is density-dependent (DD) in all cases. Unless otherwise noted, the parameters were *n*_*trips*_ = 10, 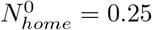 *R*_0_ = 2,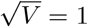, *v*_*L*_ = 1*/*3, *v*_*H*_ = 2*/*3, *η*_*H*_ = *η*_*L*_ = 1*/*2, *c*_*A*_ = *c*_*R*_ = 10.

### How does disease prevalence depend on risk perception?

Both absolute and relative risk responses exhibited complex interactions between risk perception and movement rate, *n*_*trips*_, with different forms of risk perception becoming equivalent at either extreme of movement speed (Figure 2c,d). Because solving for *I*^*^ is not analytically tractable in general, we explored these interactions further in the case where the number of trips per infectious period is very large (the “fast movement limit”) and the case where the number of trips per infectious period is very small (the “slow movement limit”). Full derivations may be found in the Supplement 4.

#### Absolute Risk Avoidance

Under absolute risk avoidance, we found that, for most parameter values and regardless of transmission mode, there was a well-defined ordering to the effectiveness of the different risk proxies: density-based perception was least effective, followed by place-based perception, and finally prevalence-based perception (which was of equivalent effectiveness as place-based perception when movement rates were very fast; Figures 2c, S1c, which show numerical results up to *n*_*trips*_ = 1000). To demonstrate this, we used the average perceived risk at equilibrium, 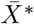, to assess differences in *I*^*^ that result from using different risk proxies. Because movement rates are monotonic functions of average perceived risk, if one risk proxy results in less perceived risk than another, that risk proxy will have a smaller effect on *I*^*^.

In the limit of fast movement, average perceived risk was the same for both place-based and prevalence-based perception and was equal to global prevalence 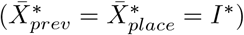. Perceived risk for density-based perception was also a function of global prevalence, but scaled by the proportional decrease in time away from home due to risk avoidance 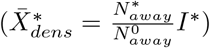. Because risk avoidance, by construction, will never cause individuals to spend more time away from home, and because density-associated risk declines when individual spend more at home, density-based perception resulted in less perceived risk than the other risk proxies and therefore had higher *I*^*^ (Figure 2c). The reason for this reduction in perceived risk is that density-based perception exhibits a negative feedback: as individuals shift to spending more time at home, the density of individuals away from home decreases, lowering perceptions of absolute risk and decreasing the perceived need to spend time at home. These results hold for both density- and frequency-dependent transmission (Figures 2c, S1c).

In the limit of slow movement, density-based perception still elicited a weaker risk response at equilibrium than did either place-based or prevalence-based perception with 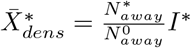 (Figure 2c, which shows numerical results down to *n*_*trips*_ = 0.001). However, unlike the fast movement limit, place-based perception differed from prevalence-based perception. Under density-dependent transmission, perception based on place was 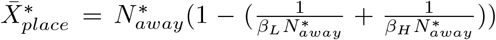 and perception based on prevalence was 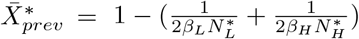. For all but the largest values of 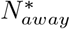, therefore, place-based perception elicited a weaker risk response than did prevalence-based perception (Figure 2c). Place-based risk perception scaled with 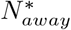 because it is equal to global prevalence 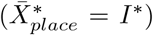, which decreases as individuals spend more time safe at home, while prevalence-based perception is the average of local prevalences, which do not comparably decrease. Similar scaling patterns are present in the frequency-dependent case, though the exact equations differ (Figure S1c; see Supplement 4).

#### Relative Risk Avoidance

Unlike absolute risk avoidance, where the ordering of different risk proxies is clear, we found that the relative effectiveness of the different risk proxies under relative risk avoidance was highly contextual. Overall, perceived risk differentials between away locations (Δ*X*), and therefore the strength of behavioral response, was directly related to spatial variation in the variable perceived as risky. Therefore, the strength of response depended heavily on the proxy used to assess risk. For example, place-based risk scaled with spatial variation in the transmission coefficients, 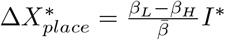 (see Figure S3c,d), and density-based risk scaled with spatial variation in density, 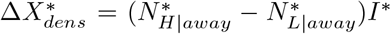, where 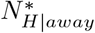 and 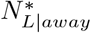 are the proportions of individuals away from home who are in the high *β* and low *β* patches, respectively (see Figure 3b,e). Moreover, prevalence-based risk showed strong differentials (and therefore elicited strong responses) in the limit of slow movement 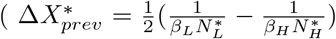 in the case of density-dependent transmission, and 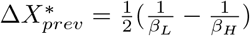 in the case of frequency-dependence). This differential disappeared in the limit of fast movement 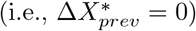 for both density- and frequency-dependent transmission, because fast movement homogenized prevalence across locations, rending prevalence ineffective as a proxy for risk.

**Figure 3:**
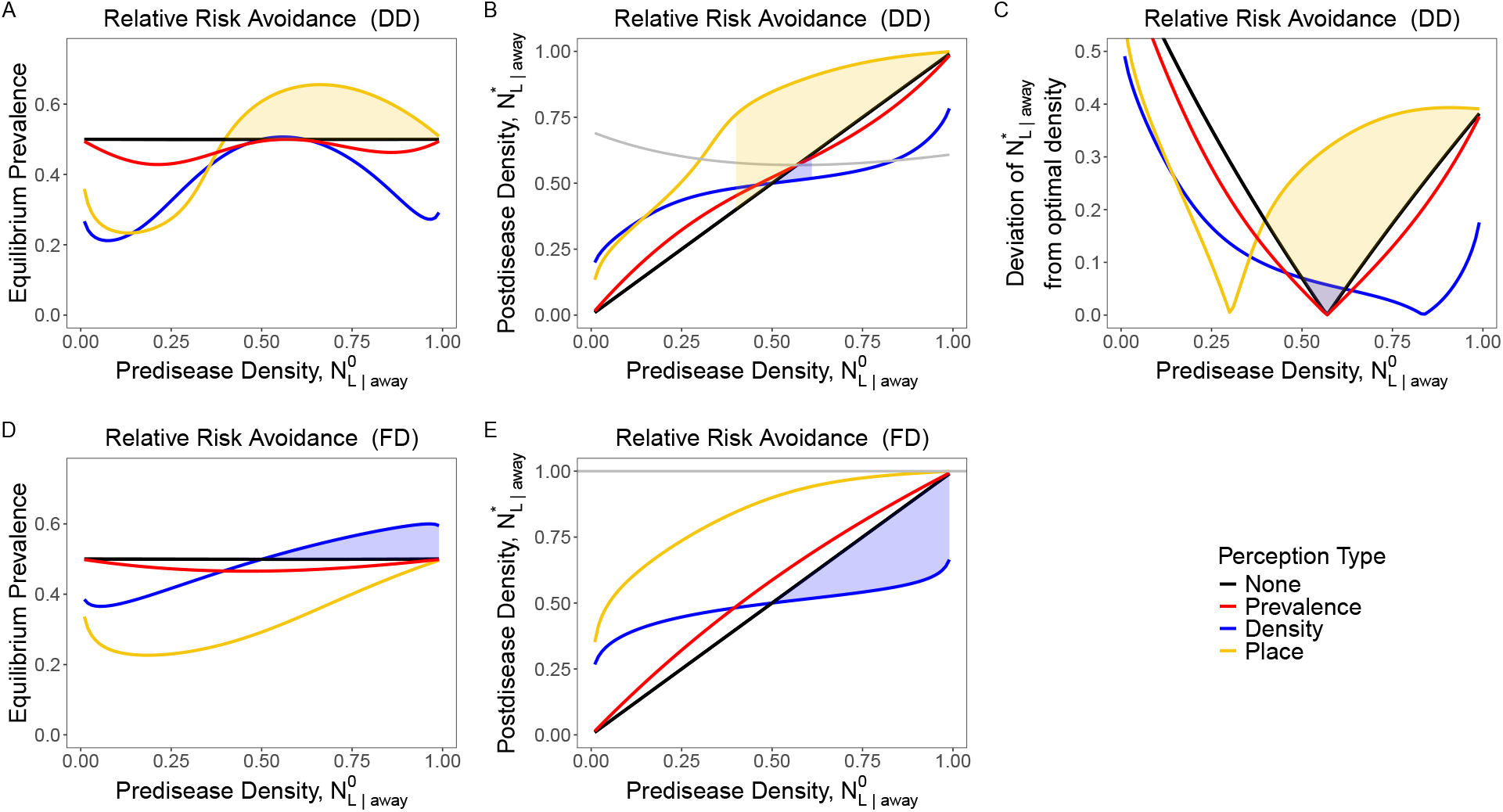
How the proportion of away-from-home individuals in the low risk patch prior to the introduction of the disease, 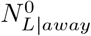, affects the equilibrium disease prevalence *I*^*^ (first column), the equilibrium proportion of away-from-home individuals in the low risk patch 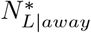 (second column), and the difference between 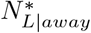 and the optimal density that minimizes equilibrium prevalence 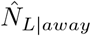 (third column) when disease transmission is density-dependent (first row) and frequency-dependent (second row). The line colors denote scenarios when risk-induced changes in travel behavior do not occur (black), individual rely on information about local prevalence (red), individuals rely on information about local host density (blue), or individuals rely on information about the transmission coefficients *β* in each location (gold). The solid grey line in B) and E) denote the optimal densities 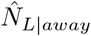under density- and frequency-dependent transmission, respectively. The shaded regions in each panel indicate values of predisease density at which a given risk proxy is misleading (i.e., results in greater equilibrium prevalence than when there are no behavioral changes). The shaded regions in panels B and C correspond to the shaded regions in panel A and the shaded region in panel E corresponds to the shaded region in panel D. Unless otherwise noted, the parameters were *n*_*trips*_ = 10, 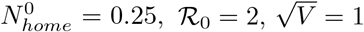, *η*_*H*_ = *η*_*L*_ = 1*/*2, *c*_*R*_ = 10.

Unlike in the case of absolute risk avoidance, when movement was slow and transmission was density-dependent, all forms of risk perception (including lack of risk perception) resulted in equivalent equilibrium values of global prevalence (Figure 2d). The fact that all risk proxies are equivalent in the limit of slow movement may be derived analytically by noting that the equilibrium prevalence has an analytical solution 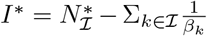, where I denotes the set of locations that sustain disease transmission (i.e., 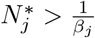). For the baseline parameters, disease transmission is sustained in all locations and therefore 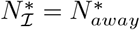, which is constant when risk avoidance is relative. Thus, no part of the equation is a function of the risk differential Δ*X*. The equivalence of relative risk proxies does not hold when transmission is frequency-dependent (Figure S1d). The nuances and implications of slow movement are further explored in Supplement 4.

### When are risk responses detrimental?

Finally, we identified circumstances under which a risk proxy can lead to detrimental behavioral responses (i.e., when risk avoidance leads to higher *I*^*^ than when disease risk is ignored). Under those circumstances, we say that such a risk proxy is “misleading”. Because the home location is safe (*β*_*home*_ = 0), absolute risk avoidance can never be detrimental and therefore no risk proxy is misleading for this type of response (see Figures 2a,c, S1a,c, S2, and S3a,b). We thus focus in this section exclusively on relative risk avoidance, where relying on the wrong risk proxy can sometimes lead to detrimental risk responses.

The circumstances for which a risk proxy was misleading depended heavily on transmission mode. Place-based perception could be detrimental if transmission was density-dependent but was never detrimental if transmission was frequency-dependent (compare the yellow lines in Figures 3a and d). By contrast, density-based perception could be detrimental when transmission was frequency-dependent, but conversely was never detrimental when transmission was density-dependent (compare the blue lines in Figures 3a and d).

To understand why place-based perception could be detrimental under density-dependent — but not frequency-dependent — transmission, consider the following heuristic based on *R*_0_ as a measure of overall disease burden. Behavioral responses alter *R*_0_ by shifting the distribution of individuals in space, because *R*_0_ is a spatial average of local *R*_0_ across away locations: *R*_0_ = *N*_*L*_ *R*_0,*L*_ + *N*_*H*_ *R*_0,*H*_, where *R*_0,*j*_ is the basic reproduction number that would result if that location were considered alone. A hypothetical optimal response 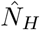 at the population level (note that 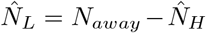) is one that minimizes R_0_: i.e., 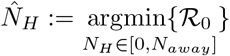. The equations for *R*_0_ in the two patch case are 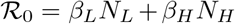 under frequency-dependent transmission, and 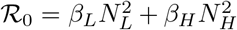 under density-dependent transmission. The population distribution that minimizes *R*_0_ in each case is 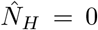 and 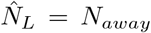 for frequency-dependent transmission, meaning all hosts are in the low transmission patch, and 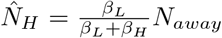and 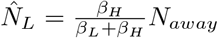 for density-dependent transmission, meaning the optimal distribution is set by the relative risk of a location as compared to the overall risk (*β*_*j*_*/*(*β*_*H*_ + *β*_*L*_)). These two optima are shown as gray lines in figures 3b,e and 4b,d (note that the *R*_0_ density-dependent optima change because is held fixed, forcing *β*_*j*_ to change as 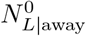 is varied).

**Figure 4.**
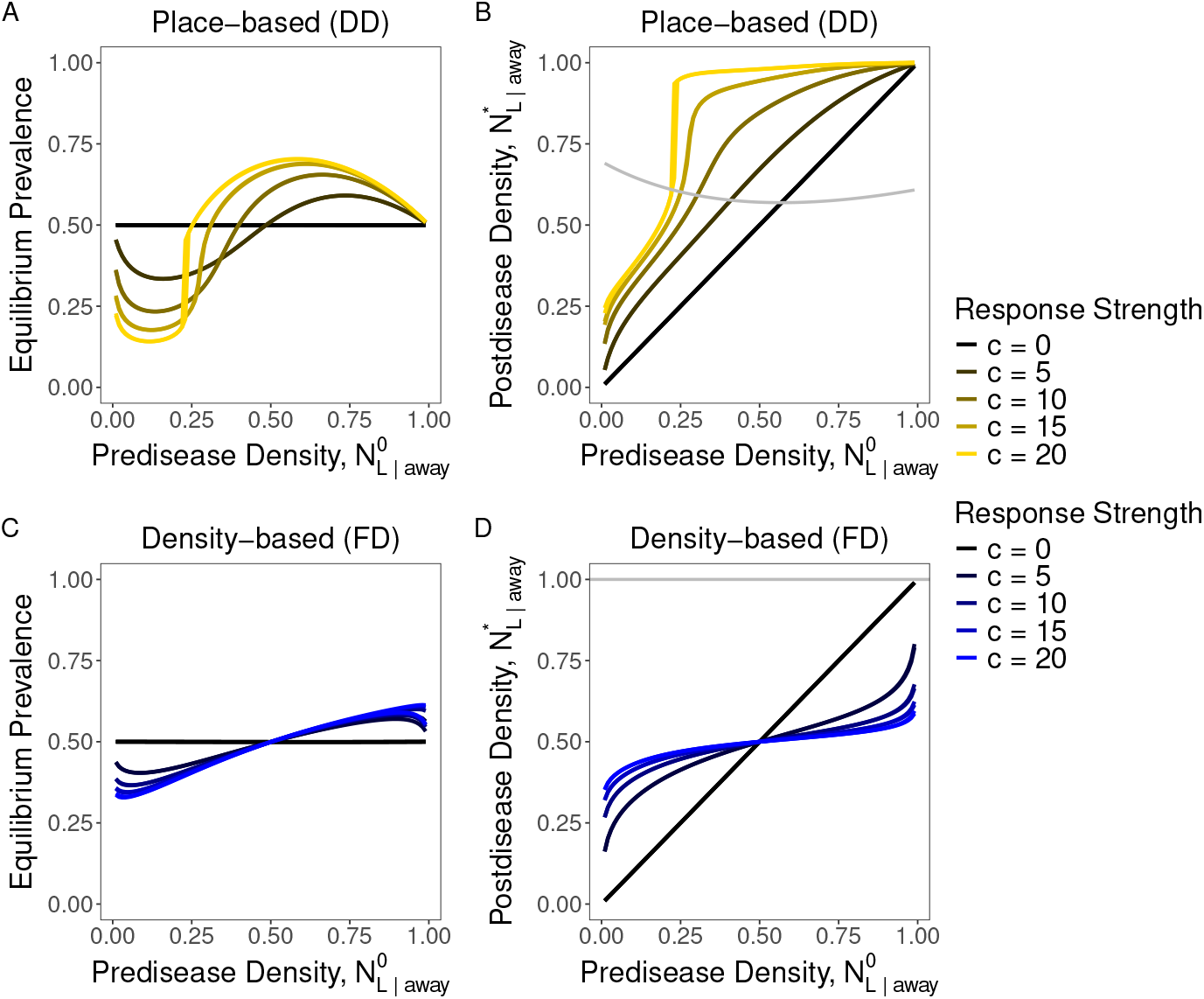
The effect of pre-disease proportion of away-from-home individuals in the low risk patch 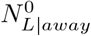 on the equilibrium disease prevalence *I*^*^ (left column) and the equilibrium proportion of away-from-home individuals in the low risk patch 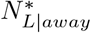 (right column) when disease transmission is density-dependent (top row) and frequency-dependent (bottom row). The solid grey lines in B) and D) denote the optimal densities 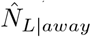 that minimize equilibrium prevalence. Unless otherwise noted, the parameters were *n*_*trips*_ = 10, 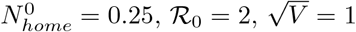 *η*_*H*_ = *η*_*L*_ = 1*/*2, *c*_*R*_ = 10.

The optimal spatial distributions indicate that the best risk avoidance strategy differs by transmission mode. Risk avoidance under frequency-dependent transmission is beneficial if it causes hosts to concentrate in the low-risk location, but is only beneficial under density-dependent transmission if it causes hosts to concentrate some, but not too much, in the low-risk location. This difference stems from the fact that high density in the less risky location can compensate for the lower transmission coefficient and lead to higher overall transmission than the ostensibly riskier location. This heuristic is approximate in general, but is exact in the case of fast movement because minimizing *R*_0_ is equivalent to minimizing *I*^*^ (Supplement 4).

Place-based perception can be detrimental when transmission is density-dependent because hosts avoid the high transmission location regardless of the host density and concentrate instead in the low transmission location. When the predisease density in the low transmission location is already high, place-based risk avoidance results in densities far in excess of what is optimal (Figure 3b), leading the spatial distribution in the absence of behavior to be closer to the optimum than the spatial distribution with place-based risk avoidance (Figure 3c). Excess avoidance of the high transmission location is facilitated by greater sensitivity to disease, *c*_*R*_, and greater initial concentration in the low disease location, 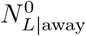 (Figure 4a)

Density-based risk avoidance can be detrimental when transmission is frequency-dependent because it equalizes the distribution of individuals across space (Figure 3e) even though the optimal response is to completely avoid the high transmission location. Thus, when most individuals are in the low transmission location pre-disease, equalizing density increases hosts in the high transmission location. By contrast, when transmission was density-dependent, the spatial smoothing caused by density avoidance brought *N*_*L*_ and *N*_*H*_ closer to their optimal values in all cases except when the initial density of individuals in the low risk location was already close to the optimum (Figure 3).

The range of 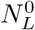 values over which place-based perception was detrimental also increased with response strength *c*_*R*_, showing an interaction between these two parameters (Figure 4a). This interaction occurred because stronger responses led 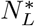 to overshoot the optimal density at smaller values of 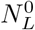 (Figure 4b). There was no such parameter interaction for the range over which density-based perception was detrimental (Figure 4c). This is because density-based perception would never lead individuals to spend more time in the higher-density location, and therefore this behavior only caused individuals to move to the high-risk location when the low-risk location had greater density. Thus while a stronger response increased the advantage or disadvantage of responding to risk, it did not change the parameter range over which the response was detrimental (Figure 4d).

## 4 Discussion

Spatial variation in infection risk means that host movement behavior plays a critical role in mediating disease transmission. When hosts are able to perceive disease risk across the landscape and alter their movement behavior accordingly, the result is a feedback loop between disease transmission and movement behavior. The complex nature of movement behavior means that individuals have many available strategies for reducing risk, while the often coarse spatial scale of disease prevalence data may force individuals to rely on other sources of information to determine local infection risk. We developed a model that decomposed two key aspects of movement behavior, risk perception and risk response, and then used this model to examine how different proxies for disease risk and different ways of avoiding risk affect the prevalence of an endemic disease. We partitioned the information used to perceive risk into three fundamental risk proxies: local disease prevalence (*I*_*j*_*/N*_*j*_), local host density (*N*_*j*_), and local transmission coefficient (*β*_*j*_; i.e., “place”). We partitioned risk avoidance into scenarios where individuals respond to risk by spending more time at home (absolute risk avoidance) and scenarios where they respond to risk by shifting their time spent away from home to relatively low risk locations (relative risk avoidance).

We found that knowing the nature of risk avoidance was crucial to ensure accurate predictions of disease transmission, as equilibrium disease prevalence was substantially lower when risk avoidance was absolute compared to relative (Figure 2). Furthermore, an absolute risk avoidance never inadvertently increased disease risk relative to ignoring risk. This contrasts markedly with relative risk avoidance, where it was possible to mistakenly perceive the riskier location as the safer one (Figure 3).

### Uncertainty in which behaviors are truly protective is critical in driving detrimental outcomes

The unequivocal benefits of absolute risk compared to relative risk in our model reflect a difference in behavioral assumptions that is often unstated in other coupled disease-behavior models. Crucial to the action of absolute risk avoidance is our assumption that home locations have no transmission. In effect, absolute risk avoidance consisted of a single decision — how frequently to travel — where one behavior — staying home — was unambiguously protective. Our absolute risk avoidance thus functions similarly to models of protective, non-spatial behaviors considered in previous studies, such as social distancing, masking, and vaccination (reviewed in Funk et al. 2010; Reitenbach et al. 2025; Verelst et al. 2016; Weston et al. 2018). Similar to these studies, using imperfect information (i.e., different risk proxies) resulted in quantitative differences in equilibrium prevalence, but never induced behaviors that increased disease risk relative to ignoring risk. The unifying feature of these studies and our absolute risk response is that the protective measure (here staying at home) is always beneficial in preventing infections.

The key feature, then, of relative risk avoidance in our model was that no single behavior was always best for reducing disease risk and the risk proxy most predictive of actual risk varied across contexts. When the information used to perceive risk was misaligned with the mode of disease transmission, individuals could exhibit detrimental risk responses even if the information was reliable in other contexts. For example, when transmission was frequency-dependent, perceiving risk based on density information led individuals to distribute themselves more evenly in space when they would have been better served by concentrating in the low risk patch (Figure 3e). Simply put, density information is irrelevant under frequency-dependent transmission. By contrast, when transmission was density-dependent, failing to use information about density or prevalence resulted in situations where individuals fueled disease transmission by excessively concentrating in the patch perceived to be low risk (Figure 3b). The exception was prevalence-based perception, which was never detrimental (though its efficacy decreased to zero in the limit of fast movement). The importance of ambiguously beneficial behaviors in driving detrimental responses is underscored by the fact that if the transmission rate at home is increased to the point where staying home can pose greater risk of infection than going out, then absolute risk responses may lead to increased disease transmission (Supplement 5; Figure S5).

These results are echoed in other models where behavioral responses to disease involve shifting movement or contacts rather than reducing them. For example, if individuals respond to disease by avoiding infected individuals but maintaining the same overall number of contacts (similar but not identical to relative risk responses in our model), this rewiring of the contact network can lead to a highly-connected cluster of susceptible individuals, which may result in cyclical outbreaks or endemic disease persistence in cases where the disease would have faded out in the absence of a behavioral response (Gross et al. 2006; Zhou et al. 2012). Similarly, if susceptible individuals respond to risk by avoiding high prevalence areas when traveling or by fleeing encounters with infected individuals, the resulting increase in movement can lead to greater connectivity across the landscape and, in turn, faster spatial disease spread (Epstein et al. 2008; Meloni et al. 2011; Nicolaides et al. 2013).

These results illustrate the interaction between two common assumptions in behavioral-epidemiological models: that hosts face a decision of whether to adopt an unambiguously protective behavior, and that prevalence (global or local) is the primary driver of perceived risk (reviewed in Funk et al. 2010; Reitenbach et al. 2025; Verelst et al. 2016). These two together can lead to overestimating the positive effects of behavioral responses by excluding possible negative correlations between perceived and actual disease risk across behaviors. Consider, for example, the behaviors of masking or not masking. While the benefits of masking relative to not masking may vary depending on environmental conditions (e.g., indoor vs outdoor), masking will never increase disease risk relative to not masking. This is to say that so long as individuals seek to reduce disease risk and the perceived benefit of masking is positively related with the actual benefit of masking, higher perceived risk will, all else equal, be positively related to higher levels of masking (though note that various factors, such as fatigue, may cause dynamic changes to the relationship between risk perception and risk response; Jørgensen et al. 2022).

In a spatial context, however, an individual’s response to disease consists of altering where they choose to spend time, and if they do not have accurate knowledge of which locations have the highest disease risk they will not have accurate knowledge of which behaviors are most protective. This allows bad information to not just weaken but reverse the relationship between perceived and actual risk associated with a particular behavior (Ruktanonchai et al. 2026). Our results showed that local prevalence, while not always an effective proxy, was never negatively correlated with actual disease transmission, while local density- and place-based information both were negative correlated with actual transmission in some circumstances. Thus, focusing only on prevalence could lead to missing possible detrimental responses, particularly in cases such as local movement where a lack of available fine-scale prevalence information may force individuals to rely on other, less consistent forms of information.

More broadly, the possibility that individuals do not accurately know which behaviors are truly protective is salient even outside the context of travel behavior. Early during the COVID-19 pandemic, for example, the relative risk of COVID transmission through aerosols or fomites was not yet clear (Chen 2021; Tellier 2022; Zheng 2020), leading to uncertainty about which environments had highest transmission and causing some people to respond with ineffective behaviors (e.g., the application of disinfecting products to produce; Geggel 2020). Additionally, as the pandemic progressed, there were major issues with misinformation, leading to people who exhibited detrimental responses to perceived risk (e.g., eschewing effective protective measures in favor of untested, ineffective, or even harmful remedies; Borges Do Nascimento et al. 2022; Van Der Linden 2022). The question of how the spread of information or disinformation about appropriate protective behaviors affects the progression of an epidemic is an important future direction (e.g., Bottemanne and Friston 2021) and is well suited to the existing “social contagion” framework, which models the spread of dual contagions (fear or disease awareness and infection) on along social networks (Fan et al. 2022; Funk et al. 2009; Granell et al. 2013; Wang et al. 2024).

Though we examined modes of risk perception based on single types of information, individuals often make decisions by integrating over multiple sources of information. While it may seem that utilizing more sources of information would improve the accuracy of risk perception, this need not necessarily be the case. For example, we found that utilizing irrelevant information on its own sometimes led to detrimental behaviors, which raises the possibility that including irrelevant information could potentially dilute the effectiveness of what would otherwise be accurate risk perception. Similarly, including accurate information in what would otherwise be inaccurate risk perception may not always be sufficient to prevent detrimental behavior. This two-fold question of when misleading risk proxies can be “fixed” by considering additional information and when otherwise accurate proxies be made misleading through extraneous information is an important direction for future research that we are actively pursuing.

Lack of accurate knowledge about the protective value of movement decisions is likely common in many circumstances. The influence of movement on population dynamics is generally known from ecological theory to be complex and highly contingent. Ecological theory dating to the 1980s shows that movement in a heterogeneous patchy landscape can boost total population size at equilibrium above the summed carrying capacity of the patches in that landscape (e.g., Holt 1985). We note that a comparable phenomenon can arise in our epidemiological system (Figures 2c,d and S1c,d). However, our model also illustrates counterexamples where increasing movement decreases prevalence (place-based and density-based perception). The details of individual movement behavior, as well as other factors such as the functional form of transmission, are all likely relevant to predicting emergent population patterns that result from individual decision making.

### Utility-based approaches: detrimental vs suboptimal behavior

We have primarily assessed the efficacy of behavioral risk responses in relation to the baseline pre-disease behavior. However, as we show in figure 3b,e, a behavioral response may also be assessed relative to the optimal response that minimizes disease risk. This approach is common for “rational epidemic” models, which treat individuals as rational agents who maximize an explicitly defined utility function. These utility functions may incorporate psychological and sociological factors such social conformity, empathy, and costs to risk-averse behavior, allowing detailed, mechanistic exploration of behavior (Eksin et al. 2017; Fenichel 2013; Fenichel et al. 2011; Morsky 2025; Morsky et al. 2023; Schnyder et al. 2025; Zhang et al. 2013). A key insight from these models is that even when individuals have perfect information, maximizing individual utility neither implies minimizing individual disease risk nor implies maximizing global utility.

Utility-based approaches are ultimately concerned with what motivates individuals. They specify the factors that influence individual decision making, explicitly modeling the costs and benefits of each choice, and allow behavior to emerge as a result. This fundamentally differs from our phenomenological approach, where we specify how individuals behave but not why they adopt a particular behavior. One may interpret the parameter values of our model in terms of the cost-benefit structure that would generate them — indeed, the very fact that individuals do not respond optimally (as seen in Figures 3b,e and 4b,d) implies constraints on the ability of individuals to minimize disease risk. For example, the proportion of time spent in each location in the absence of disease risk may be thought of as reflecting the relative utility of spending time in each location, the strengths of response *c* as reflecting the balance between negative utility associated with becoming infected and utility costs associated with risk-averse behavior, and a relative risk response as reflecting scenarios where there is a high economic or social cost to staying home (e.g., for individuals of lower socioeconomic status; Chang et al. 2021; Huang et al. 2022).

We chose to adopt a phenomenological approach over a utility-based approach for two reasons. First, it is challenging to empirically measure each component of utility (but see Bedson et al. 2021). By contrast, behavior in our model is quantified primarily in terms of movement rates and, as discussed below, is more amenable to parameterization using cell phone mobility data. Second, a phenomenological approach allowed us to study the dynamical consequences of behavior without *a priori* assumptions about why those behaviors come about. This is distinct from utility models, where the focus is on understanding the consequences of social factors influencing behavior, such as social conformity (e.g., Morsky et al. 2023; reviewed in a non-disease context by Gavrilets et al. 2024). Indeed, though we have assumed a causal link between disease risk and changes in movement behavior, the underlying model requires only correlation. Individuals who spend more time at home might do so to reduce infection risk, but they might also do so because of secondary effects of the disease — such as state-mandated restrictions or relaxed corporate work-from-home policies — that alter the underlying costs and benefits of staying home (Benzell et al. 2020; Bick et al. 2023). Thus, though we do not adopt a utility-based framework, our results may still be informative in that context, as any psychological or sociological mechanism that causes individuals to aggregate when a disease exhibits density-dependent transmission or spend less time at low-risk locations when a disease exhibits frequency-dependent transmission could prove detrimental.

The primary drawback of a phenomenological approach is that it describes the consequences of individuals relying on particular kinds of information and behaving in particular ways, but offers minimal guidance on what information individuals should use and how individuals should behave. For any particular set of circumstances we can say, for example, which risk proxy results in the lowest equilibrium prevalence. However, those circumstances themselves are a product of individual preferences and decisions (baseline movement rates, strength of behavioral response, etc…) that can’t be untangled without explicitly modeling individual decision-making. What our model can say, however, is that contextual information — how the disease spreads and how other people travel — is critical to enable good decision-making. Good information does not guarantee that everyone will make good decisions, but bad information can cause even well-meaning people to make poor decisions.

### Applications to empirical data

Though we have discussed absolute and relative risk responses as distinct, in reality individuals are likely to respond to risk by both reducing travel and avoiding high risk locations when traveling (Benzell et al. 2020; Yabe et al. 2023). This means that responses to disease risk may contain a mix of beneficial and detrimental behavior that is difficult to otherwise disentangle. In this context, one advantage to our use of a Simple Trip model of movement is that the framework lends itself to parameterization using empirical mobility data and, therefore, allows empirical patterns of mobility to be decomposed into absolute and relative components. This becomes especially important during natural disasters and major social events, where people move both due to event-related displacement and to avoid disease risk (Bengtsson et al. 2011; Finger et al. 2016). Cell phone geolocation data, for example, map hourly, fine-scale movement of individuals from small geographic units (Census Block Groups or CBGs; analogous to home locations) to specific Points of Interest (POIs; analogous to away locations), allowing time-varying estimates of the number of trips and trip duration to different POI categories (Chang et al. 2021; Yabe et al. 2023). GPS location data is also available for wildlife in the context of host movement and their influence on wildlife disease dynamics (Manlove et al. 2022; Wilber et al. 2022). Our assumption that home locations are safe reflects the scenario of the short-distance, frequent movement captured by fine-scale COVID mobility data that was used to explore trip patterns to different points of interest (Chang et al. 2021). An absence of trips is assumed to represent individuals staying in their own, separate domiciles, with lower transmission risk than if they were visiting commercial points of interest. In some circumstances, assuming that staying home minimizes infection risk is quite reasonable: This would describe individuals who live alone but have to make forays to more social environments (e.g., some student dorms) or individuals who live together but are attuned to each other’s states, so as to avoid disease transmission. At larger spatial scales, when locations represent cities or districts and trips represent longer-duration travel, the assumption of zero transmission at home becomes less plausible (e.g., Giles et al. 2021; Saucedo and Tien 2022; Wesolowski et al. 2012). While this assumption plays a critical role for absolute risk avoidance, it is unnecessary for relative risk avoidance, which reflect scenarios where seeking refuge at home is not feasible. For example, human mobility studies in dengue and malaria contexts show that even if individuals do not travel long distances, local movements and vector exposure near the home can sustain significant transmission (Stoddard et al. 2013). If anything, scenarios where home locations sustain significant transmission may be more likely to result in detrimental behavior, as reducing travel is no longer unambiguously protective (see Supplement 5;Ruktanonchai et al. 2026).

## Conclusions

Our findings highlight the important intersection between the nature of disease transmission and the nature of information used to assess disease risk in driving disease dynamics. When accurate information about disease risk is not available, or when there is widespread misinformation, individuals may use unreliable risk proxies that lead to a mismatch between perceived and actual risk. It is therefore important for disease models to incorporate imperfect information beyond just uncertainty or delays in the availability of case prevalence data and include other proxies for disease risk, as well as uncertainty or misinformation in which behaviors are most effective in reducing risk. Failing to do so may neglect detrimental behavioral responses to disease and thereby overestimate the positive effects of host behavior on disease prevention.

## Data Availability

The model scripts and output files are available on Zenodo at https://doi.org/10.5281/zenodo.21044381.

## Author Contributions

Conceptualization: all authors. Data Curation: DTC. Formal Analysis: DTC. Methodology: All authors. Funding Acquisition: NWR, NAK, OS, RDH. Software: DTC. Supervision: NAK. Visualization: DTC. Writing – Original Draft: DTC, NAK. Writing – Review and Editing: all authors.

## Supplement 1 General model formulation

In this supplement, we provide a more general formulation of the Simple Trip SIS model that allows for an arbitrary number of home and away locations, with heterogeneous movement rates among individuals from different home locations.

### Disease-free movement

Consider a population of *N* individuals divided among *K* discrete locations. Let *n*_*ij*_ denote the number of individuals that are currently in location *j* but who have homes in location *i* and let 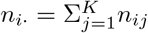 denote the total number of individuals with homes in location *i*. We define “home locations” as any location *i* such that *n*_*i*·_ *>* 0 and “away locations” as any location *i* such that *n*_*i*·_ = 0. Without loss of generality, let *K*_*home*_ be the number of home locations and *K*_*away*_ = *K* − *K*_*home*_ be the number of away locations. The total number of individuals in away location *j* is denoted 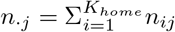 and the total number of individuals in the population is 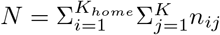. We may then normalize by the total population size so that we have the proportions *N*_*ij*_ = *n*_*ij*_*/N, N*_*i*·_ = *n*_*i*·_*/N*, and *N*_·*j*_ = *n*_·*j*_*/N* .

In the absence of disease (denoted with a superscript 0), residents of home *i* visit location *j* at a rate 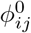. Visitation rates are composed of the total rate at which trips are taken, 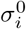, times the probability of visiting away location *j* given that a trip occurs, 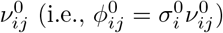. Residents of home *i* return from away location *j* at a rate 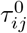, which may similarly be expressed as the total rate 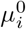 at which residents return from trips times the probability 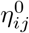 that a returning individual came from away location *j*, 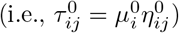). This yields the Simple Trip movement model

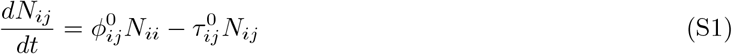

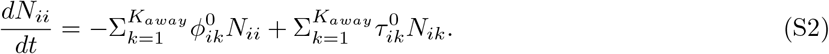

The disease-free equilibrium densities of the Simple Trip movement model are 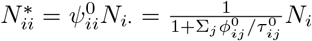 and 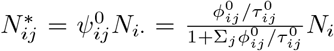., where 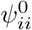 and 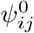 denote the diagonal and off-diagonal entries of the time allocation matrix **Ψ**^0^, which describes the average fraction of time individuals from each home location spend in each away location (Sattenspiel and Dietz 1995).

As in the main text, two individuals who occupy the same home location should not be interpreted as literally having the same domicile. The exact interpretation will depend on how home locations are defined in a particular model implementation. For example, if the model is parameterized with cell phone mobility data the individuals with the same home location may live in the same Census Block Group. In more abstract model implementations, all individuals may dwell in separate domiciles (as in the main text) and the different home locations may then represent subsets of the population with differing baseline movement behavior.

### Disease Model

The general form of the Simple Trip SIS model is written as

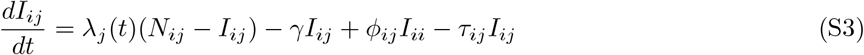

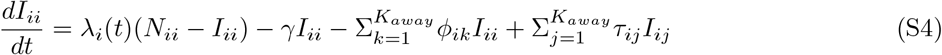

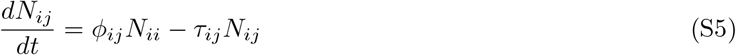

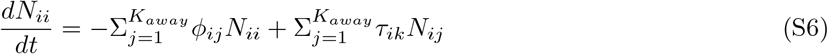

where *λ*_*j*_(*t*) is the force of infection in patch *j* at time *t* and *γ* is the rate at which infectious individuals become susceptible. We consider both density-dependent 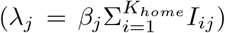 and frequency-dependent 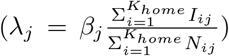 forms for the force of infection, where *β*_*j*_ is the location-specific transmission rate. Without loss of generality, we set *γ* = 1, which is equivalent to scaling time so that all rates are given in units per infectious duration (1*/γ*) and all transmission coefficients *β*_*j*_ are given in units per individual per infectious duration.

### Risk Response

We model behavioral responses to disease as changes in the visitation rates *ϕ*_*ij*_ to and return rates *τ*_*ij*_ from away locations. Baseline movement rates, 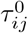 and 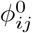, may be interpreted as the average preference for different locations in the absence of disease, and deviations from the baseline preference represent individual avoidance of (perceived) disease risk. We assume home locations have zero transmission (*β*_*i*_ = 0 for *i* ≤ *K*_*home*_), so that staying home always represents the most risk-averse behavior.

Individuals adjust their movement behavior based on perceived risk *X*_*j*_, which represents the information individuals use to assess disease risk in away location *j*. We assume that while individuals from different home locations may have differing baseline movement rates, they respond similarly to disease risk. While the main text only gives equations for changes in visitation rate, here we give equations for changes in both visitation and return rate in response to perceived disease risk.

We define “absolute risk avoidance” as the dependence of the total visitation and return rates (*σ*_*i*_ and *µ*_*i*_) on the average perceived risk 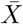 across away locations. We denote this dependence as 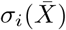 and 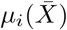), emphasizing that these rates are functions of average risk across space. We define “relative risk avoidance” as the dependence of visitation and return probabilities (*ν*_*ij*_ and *η*_*ij*_) on the perceived risk at location *j* relative to the average risk, 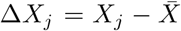 . Rates of relative movement are denoted *ν*_*ij*_(Δ*X*_*j*_) and *η*_*ij*_(Δ*X*_*j*_) to emphasize their dependence on the relative risk in location *j* compared to the spatial average.

For visitation rate, we may write

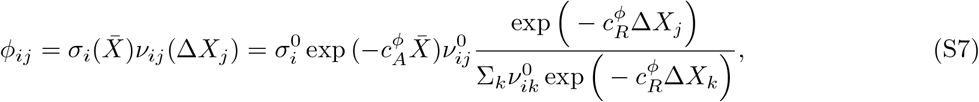

where 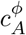 controls the absolute decrease in visitation rate with perceived disease risk and 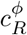 controls the relative decrease in visitation rate with perceived disease risk. The change in *ν*_*ij*_ with *X*_*j*_ is standardized so that 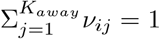.

Similarly, we may write the return rate as

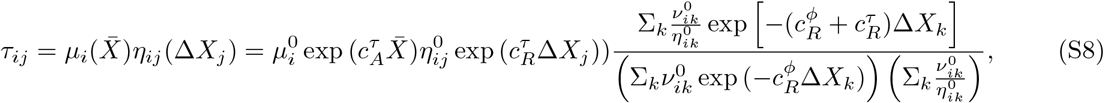

where 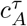 controls the absolute increase in return rate with perceived disease risk and 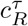 controls the relative increase in return rate with perceived disease risk. This formulation standardizes changes in *ν*_*ij*_ so that they do not alter the equilibrium fraction of time spent at home *i*, 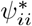. Observe that the equilibrium amount of time, for constant *X*_*j*_, spent in location *j* per unit of time spent at home *i* is

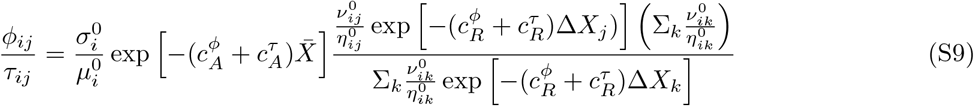

and that the equilibrium fraction of time spent at home, for constant *X*_*j*_, is

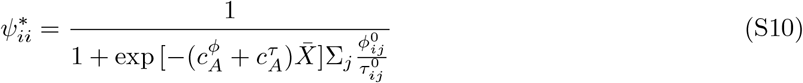

and depends only on the absolute scaling parameters *c*_1_ and *c*_3_. The equilibrium fraction of time spent away from home is then

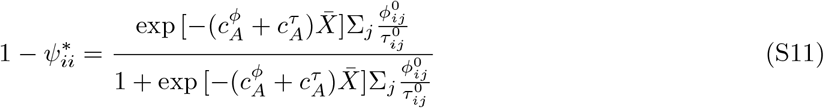

and the fraction of time spent in away location *j* is equal to

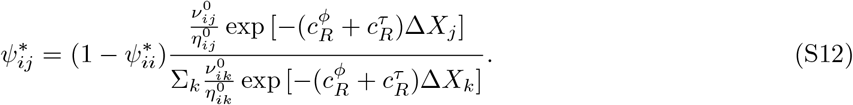

As noted above, the scaling parameters 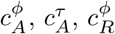, and 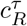 control the strength of each type of behavioral response. Setting one or more of these parameters to zero has the effect of “turning off” a given response type. Making both visitation and return rates functions of perceived disease risk allow for more complex behaviors. For example, if 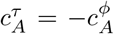 with 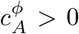 then the amount of time spent in a location does not change with perceived risk but the length of a given trip increases with 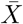 (fewer but longer trips).

## Supplement 2 Derivation of expected number of trips

The expected number of trips taken per unit time under baseline (disease-free) conditions, by the Elementary Renewal Theorem, is equal to the reciprocal of the expected time *E*[*T*] between returning home from one trip and returning home from a subsequent trip. The expected time is equal to the average time spent at home before the next trip 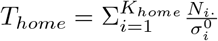 plus the average time spent per trip 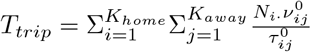 there is no variation in movement rates across individuals with different homes, the equation simplifies to 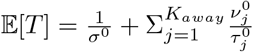. The expected number of trips may therefore be written as

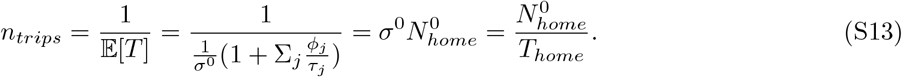

In our parameter manipulations, we vary the expected number of trips *n*_*trips*_ while holding proportion of individuals at home 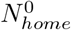 constant. However, the numerical implementation of the model requires the absolute movement rates *σ*^0^ and *µ*^0^. Consider the special case with only two away locations. The average number of trips *n*_*trips*_ and proportion of individuals at home 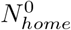 may be converted into absolute visitation *σ*^0^ and return rates *µ*^0^ through the following equations. First, equation S13 may be rearranged to solve for *σ*^0^,

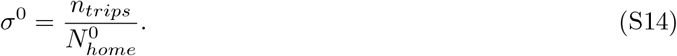

Second, note that the ratio of time spent away and time spent spent at home is equal to

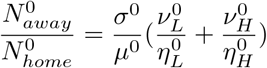

and that solving for *µ*^0^ yields

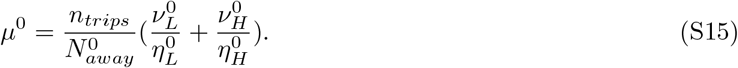

## Supplement 3 Derivation of transmission coefficients from *R*_0_ and transmission variance

In our parameter manipulations, we vary the basic reproduction number *R*_0_ independently from the spatial variance in transmission coefficients *V* . However, the numerical implementation of the model requires the transmission coefficients *β*_*H*_ and *β*_*L*_ in the high and low risk locations, respectively. In this supplement, we derive equations expressing *β*_*H*_ and *β*_*L*_ in terms of *R*_0_ and *V* .

The average of *β*_*j*_ is given by 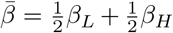 and the spatial variation of *β*_*j*_ is given by 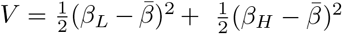. When transmission is density-dependent, 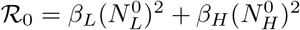. Expressing the equation for R_0_ in terms of *β*_*H*_ yields 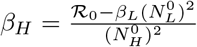, which may then be substituted into the equation for 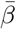 to give

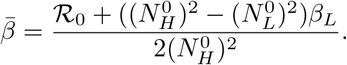

The equation for 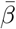 may then be substituted into the equation for *V* to yield

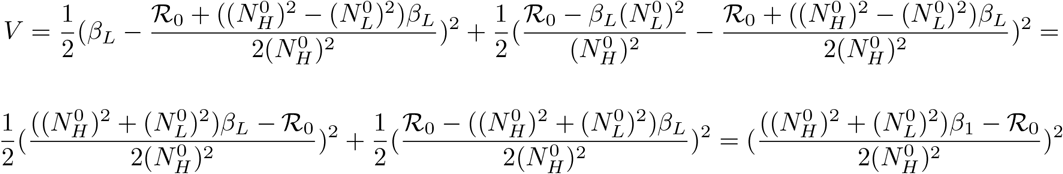

Solving for *β*_*L*_ then gives

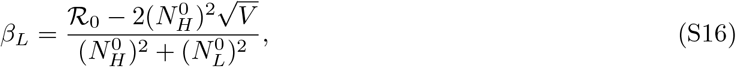

and by symmetry

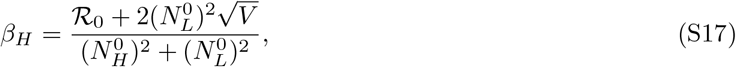

where we have taken the lesser root for the “low risk” location and the greater root for the “high risk” location.

When transmission is frequency-dependent, 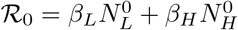. Following the same steps as above, we arrive at 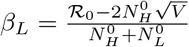 and 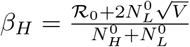.

## Supplement 4 Analytical derivations in the limits of fast and slow movement

The primary metric we use to assess the effect of risk-induced changes in movement behavior on disease transmission is the difference between equilibrium global prevalence, *I*^*^, given a particular risk response and risk proxy, and *I*^*^ in the case of risk-insensitive movement. While solving for *I*^*^ is not analytically tractable in general, useful analytic simplifications may be obtained when the number of trips per infectious period is very large (the “fast movement limit”) or when the number of trips per infectious period is very small (the “slow movement limit”). We begin our analysis using the general model formulation (see Supplement Supplement 1) before simplifying to the three patch model used in the main text.

In each of these limits, we also use the strength of behavioral response at equilibrium to assess differences in *I*^*^ that result from using different risk proxies with an absolute risk response. If two risk proxies result in the same perceived risk at equilibrium for a specific set of parameters (e.g., place-based perception and density-based perception produce the same average perceived risk, 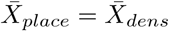, then they will produce the same equilibrium movement rates, 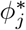 and 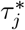, and therefore identical values of *I*^*^ (for that set of parameters). Because movement rates are monotonic functions of average perceived risk, if one risk proxy results in a weaker equilibrium response than another, then it will have a smaller effect on *I*^*^ (e.g., if 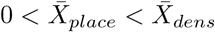, then 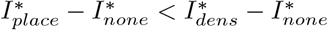).

### Fast Movement Equilibrium

In the limit of fast movement, there is time-scale separation between movement and transmission such that the spatial distribution of individuals reaches equilibrium on the timescale of transmission 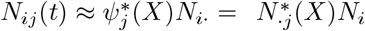, though note that when perception is density-based this equilibrium is implicit and must be solved numerically). Crucially, because movement rate does not depend on an individual’s infection status and because the rates of transmission and recovery are near zero on the timescale of movement, the distribution of susceptible and infected individuals also reaches equilibrium, with 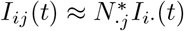 and 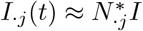. The total disease prevalence may therefore be written in the density-dependent transmission case as

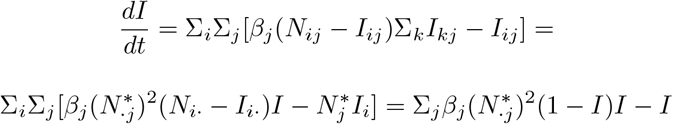

and in the frequency-dependent transmission case as

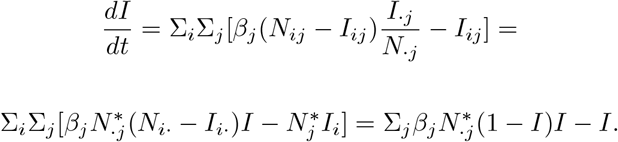

Setting 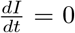 and solving for *I* yields the prevalence equilibria *I*^*^ = 0 and 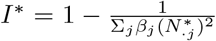 in the density-dependent case and *I*^*^ = 0 and 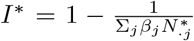 in the frequency-dependent case. The positive equilibrium is implicit and must be solved numerically, as 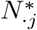 depends on *I*^*^ when risk perception is place- or prevalence-based and depends both on *I*^*^ and recursively on 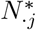 when perception is density-based. Nonetheless, we may observe from these equations that fast movement has the effect of (weighted) averaging the transmission coefficients such that 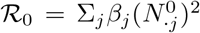 for density-dependence and 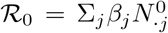 for frequency-dependence.

We may now compute the average perceived risk 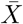 (and the strength of absolute response, all else equal) for each of the three risk proxies in the case of the three patch model presented in the main text. When perception is place-based, we have 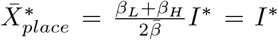. When perception is density-based we have 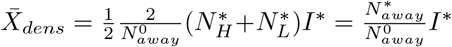. When perception is prevalence-based, 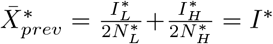. We may immediately see that place-based and prevalence-based perception are equivalent 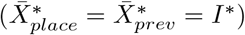, while the average perceived risk for density-based perception is discounted by a factor of 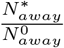, representing individual’s awareness that the number of individuals away from home (and therefore perceived risk) decreases as more individuals stay home to avoid being infected.

We may also compute the difference in perceived risk between high and low risk locations for each of the three risk proxies. This “risk differential” is defined as 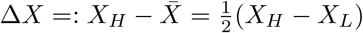 and represents the strength of relative response, all else equal. When perception is place-based, we have 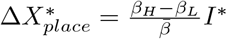.

When perception is density-based we have 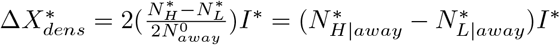 where 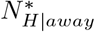 and 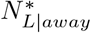 are the proportions of individuals away from home who are the high *β* and low *β* patches, respectively (note that 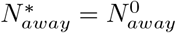 when risk response is relative). When perception is prevalence-based, 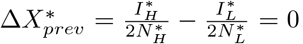, as 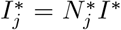. Therefore, the most evident result from these equations is that prevalence-based perception provides no information about relative disease risk and is therefore equivalent to ignoring risk altogether. These result come about because fast movement equalizes disease prevalence across space.

### Slow Movement Equilibrium

In the limit of slow movement, we may assume that each location reaches its disease equilibrium as if it were an isolated population:

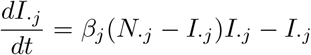

in the case of density-dependent transmission. Setting 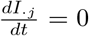 and solving for 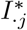 yields 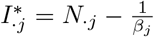 if 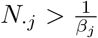 and zero otherwise.

Because transmission does not occur in home locations, individuals returning home effectively quarantine between trips when movement rates are very slow and the disease therefore cannot spread beyond the location in which it was introduced. However, in a scenario where the disease is introduced to all locations simultaneous (as is the case for our numerical simulations), we may denote the set of locations with local disease persistence as I. The global disease prevalence is then

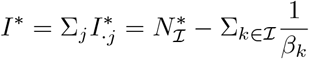

where 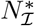 is the proportion of total individuals in locations that support disease persistence at equilibrium.

If we assume that the disease persists at equilibrium in both the high and low risk away locations, then the average perceived risk is 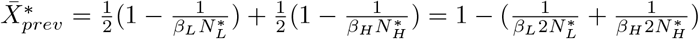 for prevalence-based perception. When perception is place-based, we have 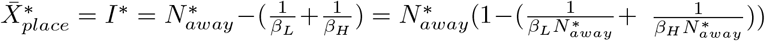. When perception is density-based, we have 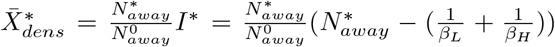. Similar to the fast-movement limit, average perceived place-based risk is equal to the equilibrium prevalence,while average perceived density-based risk is discounted by a factor of 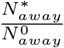. Unlike the fast movement limit, place-based perception and prevalence-based perception are not equivalent, with place-based perception tending to yield a weaker equilibrium response the more individuals are at home.

We now turn to relative risk responses. Because *β*_*j*_ do not depend on behavioral responses to disease, any two behavioral responses will produce equivalent values of *I*^*^ if they also produce equivalent values of 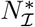. If we assume the disease persists in all away locations at equilibrium (as is the case under baseline parameters), then 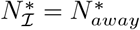. Because 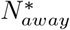 is held constant when risk avoidance is relative, *I*^*^ is insensitive to proxies used to perceive risk (including risk-insensitivity).

This counter-intuitive result holds even with an arbitrary number of away locations. Consider a comparison between the equilibrium dynamics under two risk proxies, *A* and *B*. If, for every *j*, 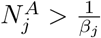 and 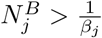 then 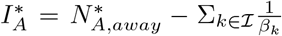 and 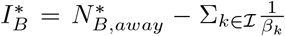. Therefore, 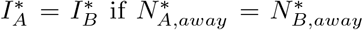. When *c*_1_ = *c*_3_ = 0 (i.e., relative risk avoidance) this is by definition always the case. We can therefore state that, in the slow movement equilibrium and when behavior does not affect the average time spent at home, different modes of risk perception produce equivalent global disease prevalence so long as the equilibrium spatial distribution of individuals given each perception variable supports disease transmission in all away locations.

Furthermore, any perception variable whose equilibrium densities do not support disease transmission in all away locations will have higher global disease prevalence than perception variables that do. Consider a change in model parameters that perturbs 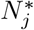 while holding 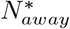 constant. Note that if 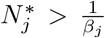 for every *j*, then 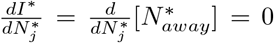. If, however, 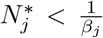 while 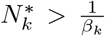 for every 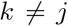, then 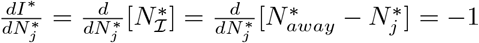. Therefore, any perception variable that results in 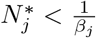 for some *j* will have greater global equilibrium disease prevalence than a perception variable that results in 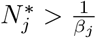 for all *j*. Moreover, for any two perception variables *A* and *B*, such that 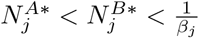 for some 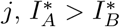.

Biologically, this result is driven by the density-dependence of transmission, which causes a one-to-one dependence of *I*_*j*_ on *N*_*j*_, and not behavioral responses to disease risk (note that the previous equations hold even when there is no behavioral response). Any decrease in *I*_*j*_ due to a shift in spatial distribution of individuals from *N*_*j*_ to *N*_*k*_ is directly offset by an increase in *I*_*k*_ so long as transmission is sustained in both locations. A shift in the distribution of individuals from an uninfected location *j* to an infected location *k* will increase *I* and a shift in the opposite direction will decrease *I* (at least until *j* reaches a critical density for sustained transmission). Because uninfected locations are precisely those in which *N*_*j*_ *<* ^1^, global prevalence is reduced when individuals become distributed more evenly across space (controlling for spatial variation in transmission coefficients).

In the case of frequency dependence, we have

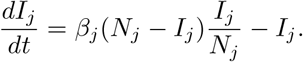

and 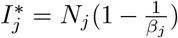 such that 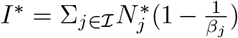, which is simply the weighted average of the equilibrium prevalences in each location. We therefore have 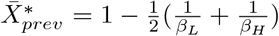 for prevalence-based perception, 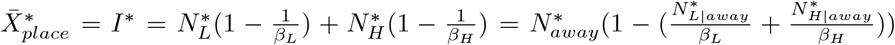 for place-based perception, and 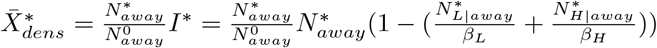 for density-based perception, so long as the disease persists in all away locations. The relationship between place-based 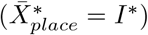 and density-based perception 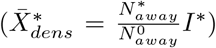 is the same as in the case of density-dependent transmission, though the equilibrium prevalence values are different. Similarly, while the equations for place-based and prevalence-based perception are different, place-based perception is still scaled by 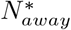, while prevalence-based perception is not.

When risk perception is relative, we have 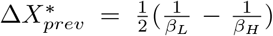 for prevalence-based perception, 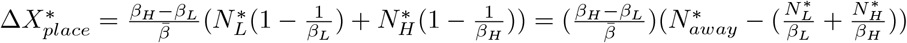 for place-based perception, and 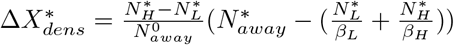 for density-based perception. The primary result to be gleaned from these equations is that, unlike with density-dependent transmission, none of the three risk proxies are equivalent.

## Supplement 5 What happens if transmission occurs at home?

In this supplement, we explore the effects of relaxing the assumption that *β*_*home*_ = 0. The main difficulty in allowing *β*_*home*_ to vary is ensuring that we make comparisons while controlling for the overall strength of transmission (i.e., holding *R*_0_ constant). We do this by splitting *R*_0_ into the sum of contributions from transmission at home and from transmission away from home 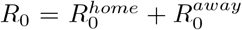. When transmission is density-dependent, 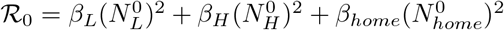. The contribution of the home location to *R*_0_ is then given by 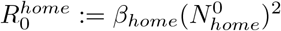. If we want vary 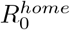 while holding *R*_0_ and *V* constant, we can simply use 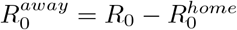 in place of *R*_0_ for equations S17 and S16. Solving for *β*_*home*_ yields 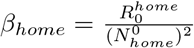. In the case of frequency-dependent transmission, the equations are similar except that 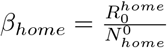

The effect of varying 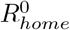 (expressed in terms of 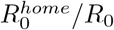) on equilibrium prevalence is shown in figure S5. In the case of relative risk avoidance, behavior does not affect the equilibrium proportion of individuals at home and so transmission in the home locations has little effect aside from weakening the ability of behavior to mitigate disease risk. In the case of absolute risk avoidance, however, transmission in the home location causes staying at home to become detrimental. The transition from beneficial to detrimental happens at much lower values of 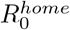 under density-dependent transmission than frequency-dependent transmission. This is due to a positive feedback between behavior and risk: when individuals reduce the rate at which they leave home, the density of individuals at home increases. This increases density-dependent transmission and, by extension, prevalence, which increases perceived risk, causing more individuals to stay home, etc…

## Supplement 6 Supplementary Figures

**Figure S1:**
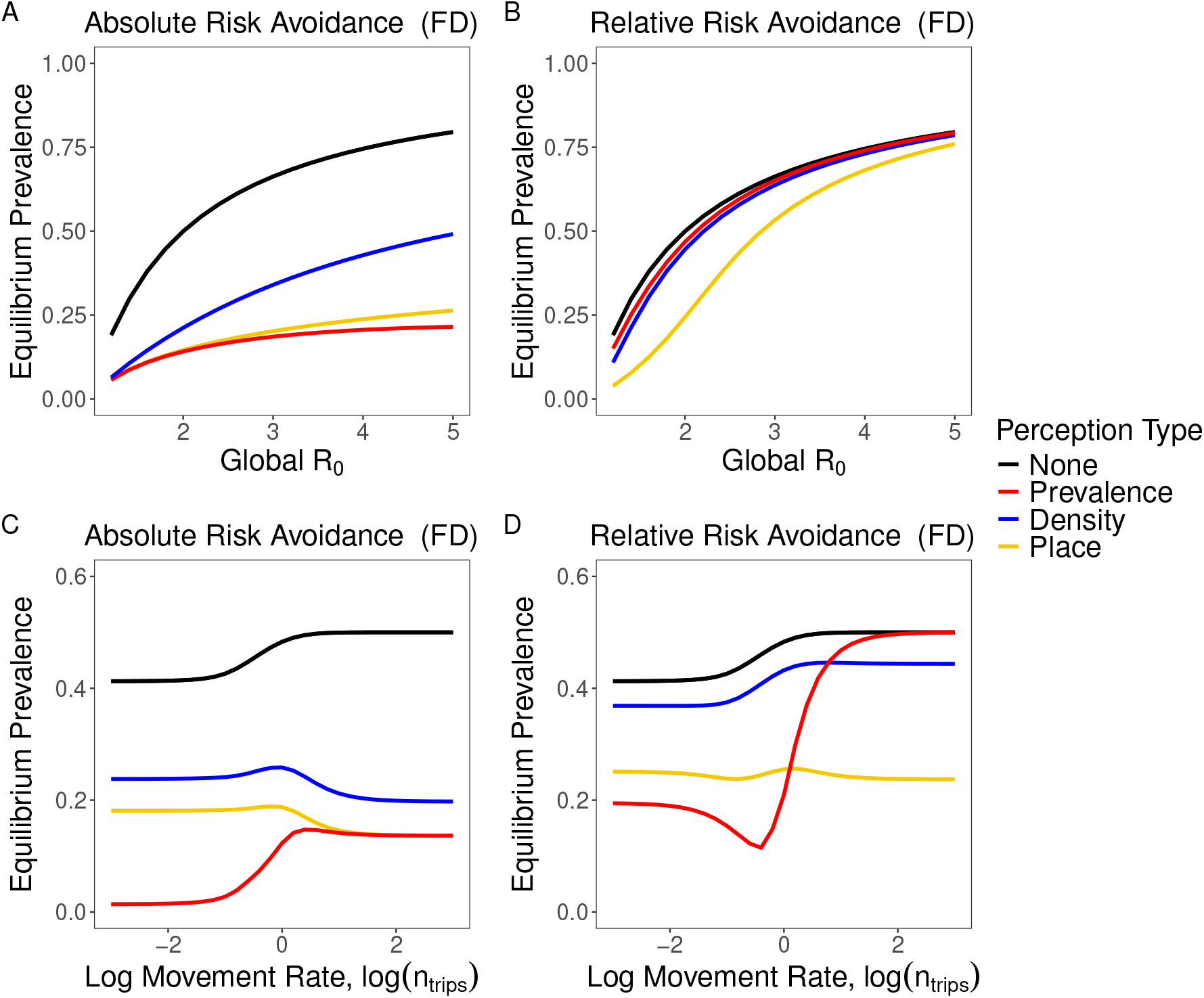
Equilibrium disease prevalence when risk-induced changes in travel behavior do not occur (black), rely on information about local prevalence (red), rely on information about local host density (blue), or rely on information about the transmission coefficients *β* in each location (gold). Top row: effect on equilibrium prevalence due to changes in *R*_0_, holding spatial variance in transmission coefficients constant. Bottom row: effect on equilibrium prevalence due to changes in log movement rate (number of trips per infectious duration). Left column: individuals respond to disease risk by increasing the proportion of time they spend at home (absolute risk response). Right column: individuals respond to disease risk by increasing the proportion of time they spend in the less risky away location, with time at home held constant. Transmission is frequency dependent (FD) in all cases. Unless otherwise noted, the parameters were 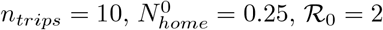, 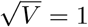, *ν*_*L*_ = 1*/*3, *ν*_*H*_ = 2*/*3, *η*_*H*_ = *η*_*L*_ = 1*/*2, *c*_*A*_ = *c*_*R*_ = 10.

**Figure S2:**
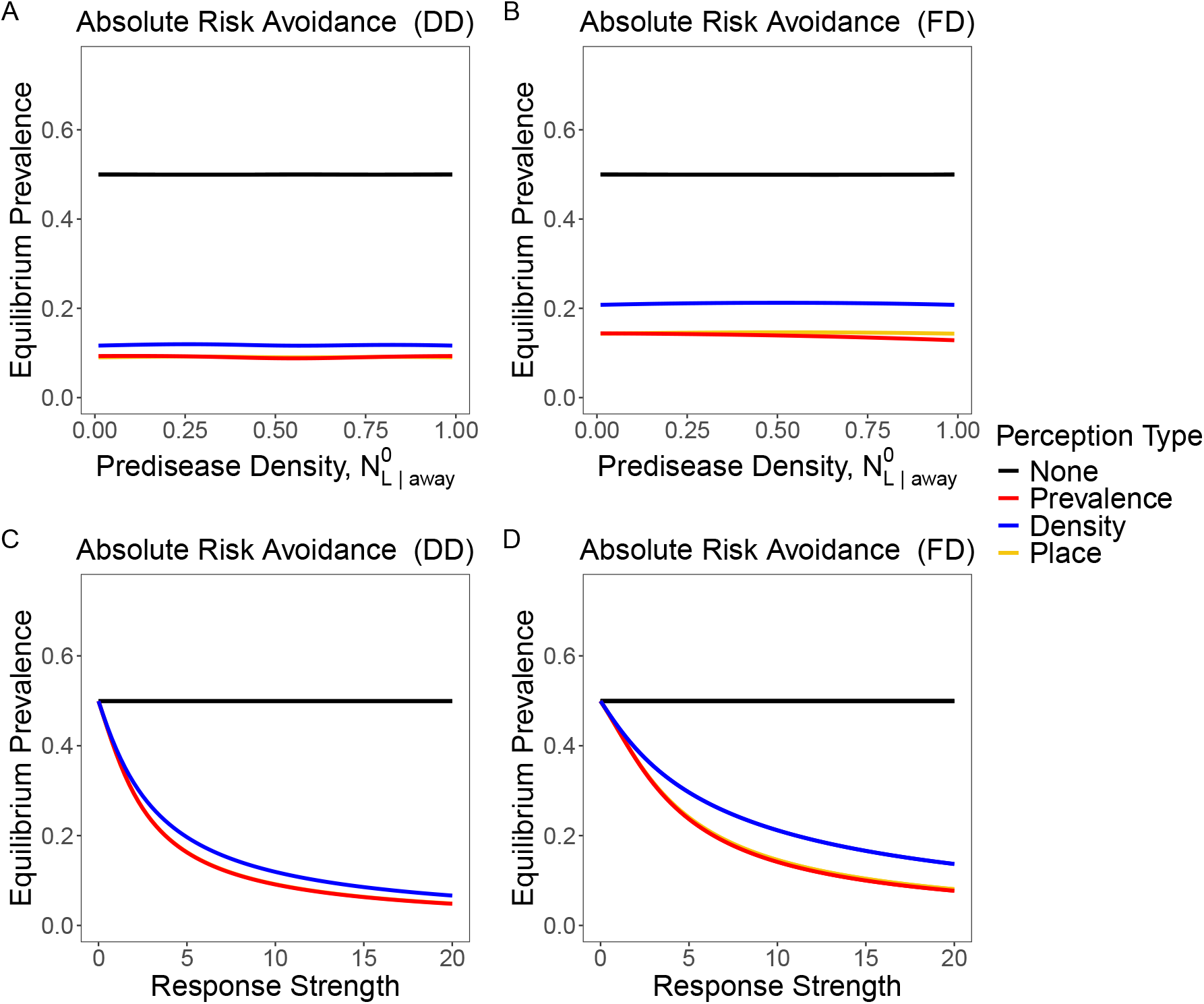
Equilibrium disease prevalence when risk-induced changes in travel behavior do not occur (black), rely on information about local prevalence (red), rely on information about local host density (blue), or rely on information about the transmission coefficients *β* in each location (gold). Top row: effect on equilibrium prevalence due to changes in the pre-disease proportion of individuals away from home that are in the low-risk location, holding *R*_0_ constant. Bottom row: the response strength *c* of movement to changes in perceived risk. Left column: disease transmission is density-dependent (DD). Right column: disease transmission is frequency dependent (FD). In all cases individuals respond to disease risk by increasing the proportion of time they spend at home (absolute risk avoidance). Unless otherwise noted, the parameters were *n*_*trips*_ = 10, 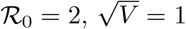, *V* = 1, *ν*_*L*_ = 1*/*3, *ν*_*H*_ = 2*/*3, *η*_*H*_ = *η*_*L*_ = 1*/*2, *c*_*A*_ = 10.

**Figure S3:**
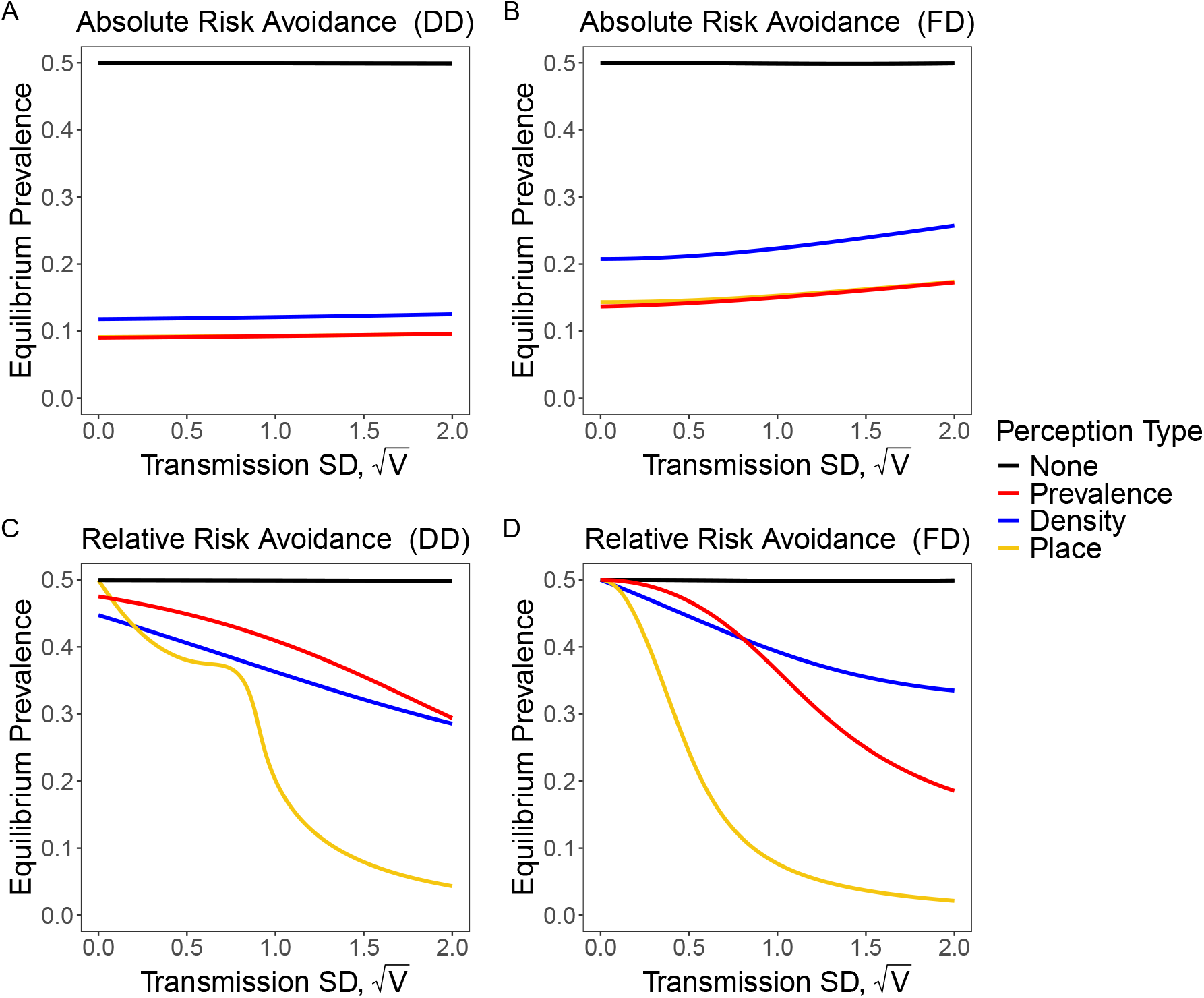
Equilibrium disease prevalence when risk-induced changes in travel behavior do not occur (black), rely on information about local prevalence (red), rely on information about local host density (blue), or rely on information about the transmission coefficients *β* in each location (gold), as a function of the spatial standard deviation of the transmission coefficients, holding *R*_0_ constant. Top row: individuals respond to disease risk by increase the proportion of time they spend at home (absolute risk avoidance). Bottom row: individuals respond to disease risk by shifting time spent in the high risk away location to the low risk away location (relative risk avoidance). Left column: disease transmission is density-dependent (DD). Right column: disease transmission is frequency dependent (FD). Unless otherwise noted, the parameters were *n*_*trips*_ = 10, 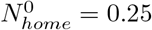, *R*_0_ = 2, *ν*_*L*_ = 1*/*3, *ν*_*H*_ = 2*/*3, *η*_*H*_ = *η*_*L*_ = 1*/*2, *c*_*A*_ = *c*_*R*_ = 10.

**Figure S4:**
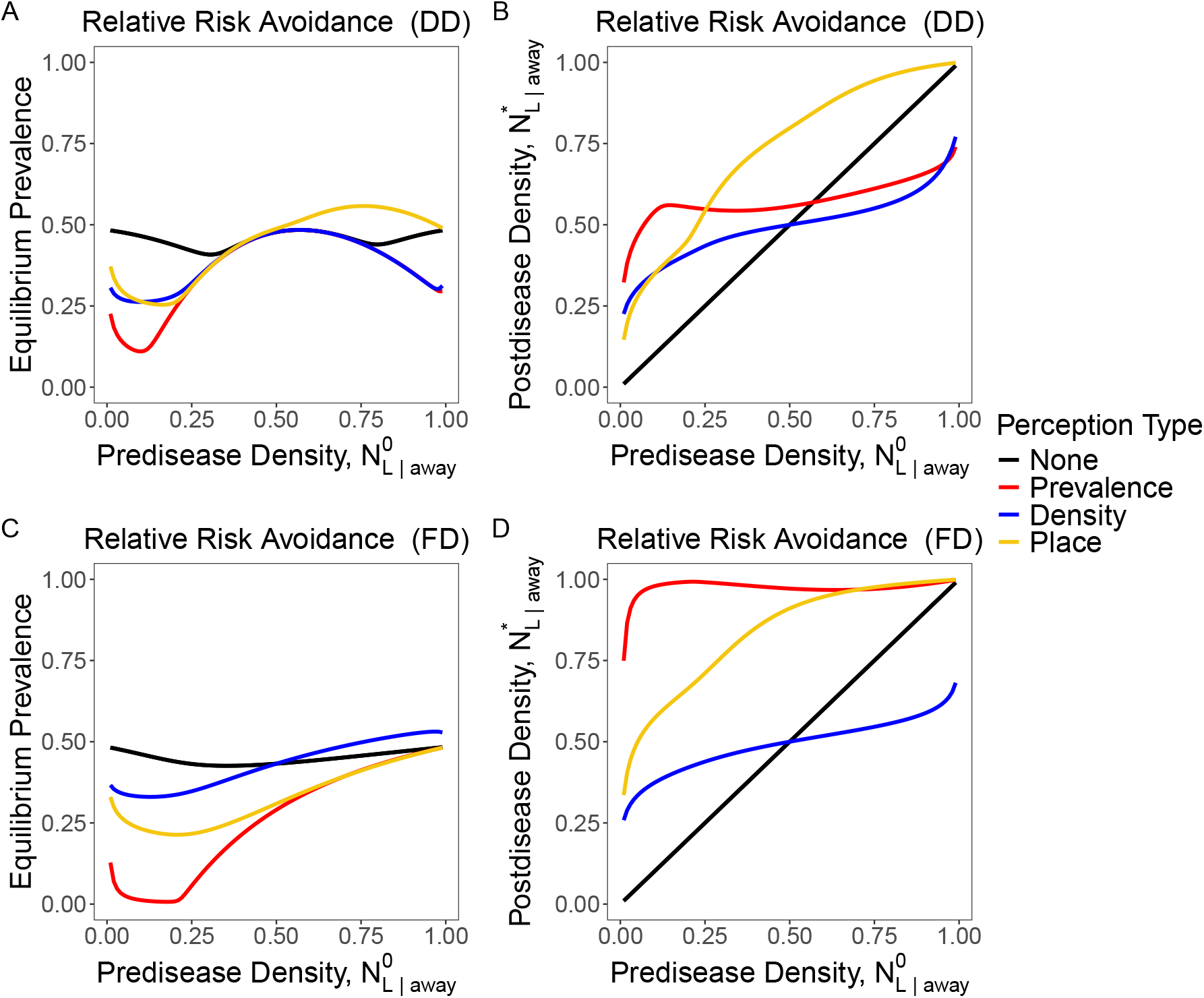
Equilibrium disease prevalence when risk-induced changes in travel behavior do not occur (black), rely on information about local prevalence (red), rely on information about local host density (blue), or rely on information about the transmission coefficients *β* in each location (gold). Top row: effect on equilibrium prevalence due to changes in the pre-disease proportion of individuals away from home that are in the low-risk location, holding *R*_0_ constant. Bottom row: the response strength *c* of movement to changes in perceived risk. Left column: disease transmission is density-dependent (DD). Right column: disease transmission is frequency dependent (FD). In all case individuals respond to disease risk by shifting their travel from more to less risky away locations (relative risk avoidance). Unless otherwise noted, the parameters were *n*_*trips*_ = 0.1, 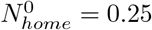, *R*_0_ = 2, 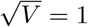, *η*_*H*_ = *η*_*L*_ = 1*/*2, *c*_*R*_ = 10.

**Figure S5:**
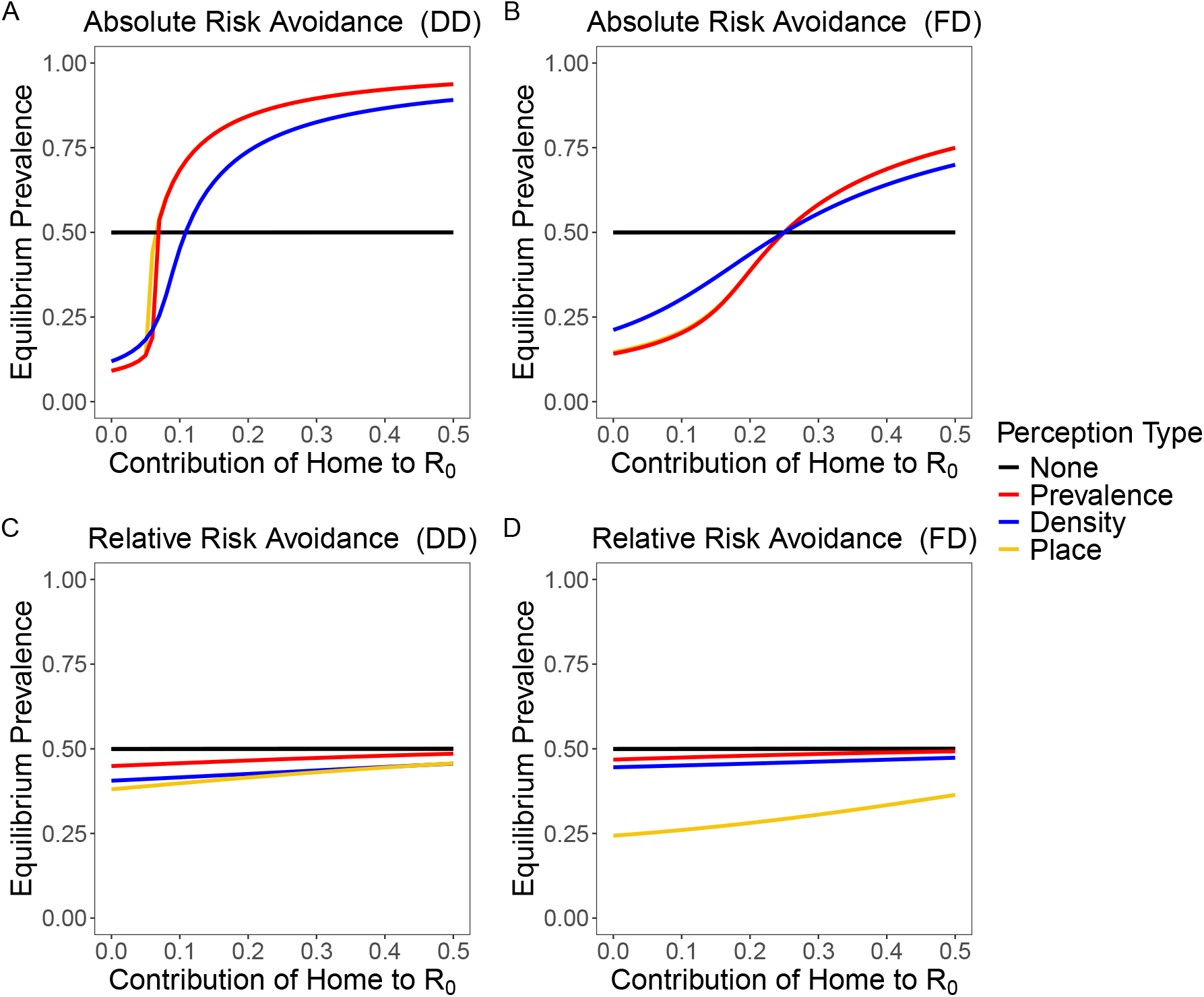
Effect of home-location transmission on equilibrium disease prevalence when risk-induced changes in travel behavior do not occur (black), rely on information about local prevalence (red), rely on information about local host density (blue), or rely on information about the transmission coefficients *β* in each location (gold). Top Row: individuals respond to disease risk by increasing the proportion of time they spend at home (absolute risk avoidance). Bottom Row: individuals respond to disease risk by shifting their travel from more to less risky away locations (relative risk avoidance). Left column: disease transmission is density-dependent (DD). Right column: disease transmission is frequency dependent (FD). *n*_*trips*_ = 10, 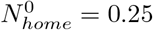, *R*_0_ = 2, *ν*_*L*_ = 1*/*3, *ν*_*H*_ = 2*/*3, *η*_*H*_ = *η*_*L*_ = 1*/*2, *c*_*A*_ = *c*_*R*_ = 10.

## References

Abdin, T. A. R., and A. H. A. Mahmoud. 2024. Lessons from the coronavirus pandemic: A review of how the disease spreads in indoor spaces. International Journal of Low-Carbon Technologies 19:90–101.

Acevedo, M. A., O. Prosper, K. Lopiano, N. Ruktanonchai, T. T. Caughlin, M. Martcheva, C. W. Osenberg, and D. L. Smith. 2015. Spatial heterogeneity, host movement and mosquito-borne disease transmission. PloS one 10:e0127552.

Agaba, G., Y. Kyrychko, and K. Blyuss. 2017. Time-delayed SIS epidemic model with population awareness. Ecological Complexity 31:50–56.

Arenas, A., W. Cota, J. Gómez-Gardeñes, S. Gómez, C. Granell, J. T. Matamalas, D. Soriano-Paños, and B. Steinegger. 2020. Modeling the spatiotemporal epidemic spreading of covid-19 and the impact of mobility and social distancing interventions. Physical Review X 10:041055.

Arthur, R. F., J. H. Jones, M. H. Bonds, Y. Ram, and M. W. Feldman. 2021. Adaptive social contact rates induce complex dynamics during epidemics. PLoS Computational Biology 17:e1008639.

Banerjee, M., V. Volpert, P. Manfredi, and A. d’Onofrio. 2024. Behavior-induced phase transitions with far from equilibrium patterning in a SIS epidemic model: Global vs non-local feedback. Physica D: Nonlinear Phenomena 469:134316.

Bedson, J., L. A. Skrip, D. Pedi, S. Abramowitz, S. Carter, M. F. Jalloh, S. Funk, N. Gobat, T. Giles-Vernick, G. Chowell, J. R. de Almeida, R. Elessawi, S. V. Scarpino, R. A. Hammond, S. Briand, J. M. Epstein, L. Hébert-Dufresne, and B. M. Althouse. 2021. A review and agenda for integrated disease models including social and behavioural factors. Nature Human Behaviour 5:834–846.

Bengtsson, L., X. Lu, A. Thorson, R. Garfield, and J. Von Schreeb. 2011. Improved response to disasters and outbreaks by tracking population movements with mobile phone network data: A post-earthquake geospatial study in haiti. PLoS medicine 8:e1001083.

Benzell, S. G., A. Collis, and C. Nicolaides. 2020. Rationing social contact during the COVID-19 pandemic: Transmission risk and social benefits of US locations. Proceedings of the National Academy of Sciences 117:14642–14644.

Bick, A., A. Blandin, and K. Mertens. 2023. Work from Home before and after the COVID-19 Outbreak. American Economic Journal: Macroeconomics 15:1–39.

Borges Do Nascimento, I. J., A. Beatriz Pizarro, J. Almeida, N. Azzopardi-Muscat, M. André Gonçalves, M. Björklund, and D. Novillo-Ortiz. 2022. Infodemics and health misinformation: A systematic review of reviews. Bulletin of the World Health Organization 100:544–561.

Bottemanne, H., and K. J. Friston. 2021. An active inference account of protective behaviours during the covid-19 pandemic. Cognitive, Affective, & Behavioral Neuroscience 21:1117–1129.

Chang, S., E. Pierson, P. W. Koh, J. Gerardin, B. Redbird, D. Grusky, and J. Leskovec. 2021. Mobility network models of COVID-19 explain inequities and inform reopening. Nature 589:82–87.

Changruenngam, S., D. J. Bicout, and C. Modchang. 2020. How the individual human mobility spatiotemporally shapes the disease transmission dynamics. Scientific Reports 10:11325.

Chen, F. H. 2009. Modeling the effect of information quality on risk behavior change and the transmission of infectious diseases. Mathematical Biosciences 217:125–133.

Chen, J., A. Marathe, and M. Marathe. 2018. Feedback between behavioral adaptations and disease dynamics. Scientific Reports 8:12452.

Chen, T. 2021. Fomites and the covid-19 pandemic: An evidence review on its role in viral transmission. National Collaborating Centre for Environmental Health: Vancouver, BC, Canada pages 1–24.

Cheng, T., and J. Wu. 2025. Recurrent patterns of disease spread post the acute phase of a pandemic: Insights from a coupled system of a differential equation for disease transmission and a delayed algebraic equation for behavioral adaptation. Mathematical Biosciences 387:109480.

Cheng, T., and X. Zou. 2024. Modelling the impact of precaution on disease dynamics and its evolution. Journal of Mathematical Biology 89:1.

Citron, D. T., C. A. Guerra, A. J. Dolgert, S. L. Wu, J. M. Henry H. M. Sánchez C., and D. L. Smith. 2021. Comparing metapopulation dynamics of infectious diseases under different models of human movement. Proceedings of the National Academy of Sciences 118:e2007488118.

Costello, F., P. Watts, and R. Howe. 2023. A model of behavioural response to risk accurately predicts the statistical distribution of COVID-19 infection and reproduction numbers. Scientific Reports 13:2435.

Dashtbali, M., and M. Mirzaie. 2021. A compartmental model that predicts the effect of social distancing and vaccination on controlling covid-19. Scientific Reports 11:8191.

Daversa, D., A. Fenton, A. Dell, T. Garner, and A. Manica. 2017. Infections on the move: How transient phases of host movement influence disease spread. Proceedings of the Royal Society B: Biological Sciences 284:20171807.

Demirel, G., E. Barter, and T. Gross. 2017. Dynamics of epidemic diseases on a growing adaptive network. Scientific Reports 7:42352.

DeWitt, M. E., B. R. Bellotti, and N. Kortessis. 2024. Time is of the essence: Effectiveness of dairy farm control of h5n1 is limited by fast spread. bioRxiv .

Dolfi, A. C., K. Kausrud, K. Rysava, C. Champagne, Y.-H. Huang, Z. R. Barandongo, and W. C. Turner. 2024. Season of death, pathogen persistence and wildlife behaviour alter number of anthrax secondary infections from environmental reservoirs. Proceedings of the Royal Society B: Biological Sciences 291:20232568.

Dougherty, E. R., D. P. Seidel, J. K. Blackburn, W. C. Turner, and W. M. Getz. 2022. A framework for integrating inferred movement behavior into disease risk models. Movement Ecology 10:31.

Eksin, C., K. Paarporn, and J. S. Weitz. 2019. Systematic biases in disease forecasting – The role of behavior change. Epidemics 27:96–105.

Eksin, C., J. S. Shamma, and J. S. Weitz. 2017. Disease dynamics in a stochastic network game: A little empathy goes a long way in averting outbreaks. Scientific Reports 7:44122.

Epstein, J. M., J. Parker, D. Cummings, and R. A. Hammond. 2008. Coupled contagion dynamics of fear and disease: Mathematical and computational explorations. PLoS ONE 3:e3955.

Ezenwa, V. O. 2004. Selective defecation and selective foraging: Antiparasite behavior in wild ungulates? Ethology 110:851–862.

Ezenwa, V. O., E. A. Archie, M. E. Craft, D. M. Hawley, L. B. Martin, J. Moore, and L. White. 2016. Host behaviour–parasite feedback: An essential link between animal behaviour and disease ecology. Proceedings of the Royal Society B: Biological Sciences 283:20153078.

Fair, K. R., V. A. Karatayev, M. Anand, and C. T. Bauch. 2024. Impact of population behavioural responses on the critical community size of infectious diseases. Theoretical Ecology 17:269–280.

Fan, J., Q. Yin, C. Xia, and M. Perc. 2022. Epidemics on multilayer simplicial complexes. Proceedings of the Royal Society A: Mathematical, Physical and Engineering Sciences 478:20220059.

Fenichel, E. P. 2013. Economic considerations for social distancing and behavioral based policies during an epidemic. Journal of Health Economics 32:440–451.

Fenichel, E. P., C. Castillo-Chavez, M. G. Ceddia, G. Chowell, P. A. G. Parra, G. J. Hickling, G. Holloway, R. Horan, B. Morin, C. Perrings, M. Springborn, L. Velazquez, and C. Villalobos. 2011. Adaptive human behavior in epidemiological models. Proceedings of the National Academy of Sciences 108:6306–6311.

Finger, F., T. Genolet, L. Mari, G. C. de Magny, N. M. Manga, A. Rinaldo, and E. Bertuzzo. 2016. Mobile phone data highlights the role of mass gatherings in the spreading of cholera outbreaks. Proceedings of the National Academy of Sciences 113:6421–6426.

Funk, S., E. Gilad, C. Watkins, and V. A. A. Jansen. 2009. The spread of awareness and its impact on epidemic outbreaks. Proceedings of the National Academy of Sciences 106:6872–6877.

Funk, S., M. Salathé, and V. A. A. Jansen. 2010. Modelling the influence of human behaviour on the spread of infectious diseases: A review. Journal of The Royal Society Interface 7:1247–1256.

Gavrilets, S., D. Tverskoi, and A. Sánchez. 2024. Modelling social norms: An integration of the norm-utility approach with beliefs dynamics. Philosophical Transactions of the Royal Society B: Biological Sciences 379:20230027.

Geggel, L. 2020. Viral video advises washing fruit and vegetables with soap. Here’s why that’s a bad idea. Live Science.

Gibson, A. K., and C. R. Amoroso. 2022. Evolution and Ecology of Parasite Avoidance. Annual Review of Ecology, Evolution, and Systematics 53:47–67.

Giles, J. R., D. A. Cummings, B. T. Grenfell, A. J. Tatem, E. z. Erbach-Schoenberg, C. J. E. Metcalf, and A. Wesolowski. 2021. Trip duration drives shift in travel network structure with implications for the predictability of spatial disease spread. PLoS computational biology 17:e1009127.

Giles, J. R., E. Zu Erbach-Schoenberg, A. J. Tatem, L. Gardner, O. N. Bjørnstad, C. J. E. Metcalf, and A. Wesolowski. 2020. The duration of travel impacts the spatial dynamics of infectious diseases. Proceedings of the National Academy of Sciences 117:22572–22579.

Granell, C., S. Gómez, and A. Arenas. 2013. Dynamical interplay between awareness and epidemic spreading in multiplex networks. Physical Review Letters 111:128701.

Gross, T., C. J. D. D’Lima, and B. Blasius. 2006. Epidemic dynamics on an adaptive network. Physical Review Letters 96:208701.

Hagenaars, T., C. Donnelly, and N. Ferguson. 2004. Spatial heterogeneity and the persistence of infectious diseases. Journal of theoretical biology 229:349–359.

Hamilton, A., F. Haghpanah, A. Tulchinsky, N. Kipshidze, S. Poleon, G. Lin, H. Du, L. Gardner, and E. Klein. 2024. Incorporating endogenous human behavior in models of COVID-19 transmission: A systematic scoping review. Dialogues in Health 4:100179.

Hawley, D. M., A. K. Gibson, A. K. Townsend, M. E. Craft, and J. F. Stephenson. 2021. Bidirectional interactions between host social behaviour and parasites arise through ecological and evolutionary processes. Parasitology 148:274–288.

Holt, R. D. 1985. Population dynamics in two-patch environments: some anomalous consequences of an optimal habitat distribution. Theoretical population biology 28:181–208.

Hoverman, J. T., and C. L. Searle. 2016. Behavioural influences on disease risk: Implications for conservation and management. Animal Behaviour 120:263–271.

Huang, X., J. Lu, S. Gao, S. Wang, Z. Liu, and H. Wei. 2022. Staying at home is a privilege: Evidence from fine-grained mobile phone location data in the united states during the COVID-19 pandemic. Annals of the American Association of Geographers 112:286–305.

Jørgensen, F., A. Bor, M. S. Rasmussen, M. F. Lindholt, and M. B. Petersen. 2022. Pandemic fatigue fueled political discontent during the COVID-19 pandemic. Proceedings of the National Academy of Sciences 119:e2201266119.

Koher, A., F. Jørgensen, M. B. Petersen, and S. Lehmann. 2023. Epidemic modelling of monitoring public behavior using surveys during pandemic-induced lockdowns. Communications Medicine 3:80.

Kumar, V., C. T. Bauch, and S. Bhattacharyya. 2024. A game theoretic complex network model to estimate the epidemic threshold under individual vaccination behaviour and adaptive social connections. Scientific Reports 14:29148.

LaDeau, S. L., B. F. Allan, P. T. Leisnham, and M. Z. Levy. 2015. The ecological foundations of transmission potential and vector-borne disease in urban landscapes. Functional ecology 29:889–901.

LeJeune, L., N. Ghaffarzadegan, L. M. Childs, and O. Saucedo. 2024. Mathematical analysis of simple behavioral epidemic models. Mathematical Biosciences 375:109250.

LeJeune, L., N. Ghaffarzadegan, L. M. Childs, and O. Saucedo. 2025a. Formulating human risk response in epidemic models: Exogenous vs endogenous approaches. European Journal of Operational Research 324:246–258.

LeJeune, L., O. Saucedo, L. M. Childs, and N. Ghaffarzadegan. 2025b. The behavioral spillover effect: Modeling behavioral interdependencies in multi-pathogen dynamics. arXiv preprint arXiv:2510.19153 .

Leung, N. H. L. 2021. Transmissibility and transmission of respiratory viruses. Nature Reviews Microbiology 19:528–545.

Li, Y., H. Qian, J. Hang, X. Chen, P. Cheng, H. Ling, S. Wang, P. Liang, J. Li, S. Xiao, et al. 2021. Probable airborne transmission of SARS-CoV-2 in a poorly ventilated restaurant. Building and Environment 196:107788.

Lloyd, A. L., and R. M. May. 1996. Spatial heterogeneity in epidemic models. Journal of Theoretical Biology 179:1–11.

Lobov, S. A., A. I. Zharinov, E. S. Berdnikova, D. P. Kurganov, and V. B. Kazantsev. 2024. Information feedback provokes multi-peak dynamics in the modern pandemic spreading. Nonlinear Dynamics 112:14677–14686.

Machens, A., F. Gesualdo, C. Rizzo, A. E. Tozzi, A. Barrat, and C. Cattuto. 2013. An infectious disease model on empirical networks of human contact: Bridging the gap between dynamic network data and contact matrices. BMC Infectious Diseases 13:185.

Maharaj, S., and A. Kleczkowski. 2012. Controlling epidemic spread by social distancing: Do it well or not at all. BMC public health 12:679.

Mahmud, M. S., S. Eshun, B. Espinoza, and C. Kadelka. 2025. Adaptive human behavior and delays in information availability autonomously modulate epidemic waves. PNAS nexus 4:pgaf145.

Manlove, K., M. Wilber, L. White, G. Bastille-Rousseau, A. Yang, M. L. J. Gilbertson, M. E. Craft, P. C. Cross, G. Wittemyer, and K. M. Pepin. 2022. Defining an epidemiological landscape that connects movement ecology to pathogen transmission and pace-of-life. Ecology Letters 25:1760–1782.

McCallum, H., A. Fenton, P. J. Hudson, B. Lee, B. Levick, R. Norman, S. E. Perkins, M. Viney, A. J. Wilson, and J. Lello. 2017. Breaking beta: Deconstructing the parasite transmission function. Philosophical Transactions of the Royal Society B: Biological Sciences 372:20160084.

Meloni, S., N. Perra, A. Arenas, S. Gómez, Y. Moreno, and A. Vespignani. 2011. Modeling human mobility responses to the large-scale spreading of infectious diseases. Scientific Reports 1:62.

Mistry, D., M. Litvinova, A. Pastore Y Piontti, M. Chinazzi, L. Fumanelli, M. F. C. Gomes, S. A. Haque, Q.-H. Liu, K. Mu, X. Xiong, M. E. Halloran, I. M. Longini, S. Merler, M. Ajelli, and A. Vespignani. 2021. Inferring high-resolution human mixing patterns for disease modeling. Nature Communications 12:323.

Morsky, B. 2025. Vaccination and collective action under social norms. Bulletin of Mathematical Biology 87:55.

Morsky, B., F. Magpantay, T. Day, and E. Akçay. 2023. The impact of threshold decision mechanisms of collective behavior on disease spread. Proceedings of the National Academy of Sciences 120:e2221479120.

Müller, S. A., M. Balmer, W. Charlton, R. Ewert, A. Neumann, C. Rakow, T. Schlenther, and K. Nagel. 2021. Predicting the effects of COVID-19 related interventions in urban settings by combining activity-based modelling, agent-based simulation, and mobile phone data. PLOS ONE 16:e0259037.

Nathan, R., W. M. Getz, E. Revilla, M. Holyoak, R. Kadmon, D. Saltz, and P. E. Smouse. 2008. A movement ecology paradigm for unifying organismal movement research. Proceedings of the National Academy of Sciences 105:19052–19059.

Nicolaides, C., L. Cueto-Felgueroso, and R. Juanes. 2013. The price of anarchy in mobility-driven contagion dynamics. Journal of The Royal Society Interface 10:20130495.

Osi, A., and N. Ghaffarzadegan. 2024. Parameter estimation in behavioral epidemic models with endogenous societal risk-response. PLOS Computational Biology 20:e1011992.

Perra, N. 2021. Non-pharmaceutical interventions during the COVID-19 pandemic: A review. Physics Reports 913:1–52.

Perra, N., D. Balcan, B. Gonçalves, and A. Vespignani. 2011. Towards a Characterization of Behavior-Disease Models. PLOS ONE 6:e23084.

Qiu, Z., B. Espinoza, V. V. Vasconcelos, C. Chen, S. M. Constantino, S. A. Crabtree, L. Yang, A. Vullikanti, J. Chen, J. Weibull, K. Basu, A. Dixit, S. A. Levin, and M. V. Marathe. 2022. Understanding the coevolution of mask wearing and epidemics: A network perspective. Proceedings of the National Academy of Sciences 119:e2123355119.

Rahmandad, H., R. Xu, and N. Ghaffarzadegan. 2022. Enhancing long-term forecasting: Learning from COVID-19 models. PLOS Computational Biology 18:e1010100.

Reitenbach, A., F. Sartori, S. Banisch, A. Golovin, A. Calero Valdez, M. Kretzschmar, V. Priesemann, and M. Mäs. 2025. Coupled infectious disease and behavior dynamics. A review of model assumptions. Reports on Progress in Physics 88:016601.

Rodríguez, D. J., and L. Torres-Sorando. 2001. Models of infectious diseases in spatially heterogeneous environments. Bulletin of Mathematical Biology 63:547–571.

Romeo-Aznar, V., L. Picinini Freitas, O. Gonçalves Cruz, A. A. King, and M. Pascual. 2022. Fine-scale heterogeneity in population density predicts wave dynamics in dengue epidemics. Nature Communications 13:996.

Ruktanonchai, N. W., N. Kortessis, D. T. Clement, O. Saucedo, E. Cleary, S. Lai, S. Gao, and R. D. Holt. 2026. Accounting for a typology of travel behavior in modeling infectious disease transmission, detection, and community response. Scientific Reports .

Ryan, M., E. Brindal, and R. I. Hickson. 2025. Behaviour and infection feedback loops inform early-stage behaviour emergence and the efficacy of interventions. Mathematics in Medical and Life Sciences 2:2452444.

Ryan, M., E. Brindal, M. Roberts, and R. I. Hickson. 2024. A behaviour and disease transmission model: incorporating the Health Belief Model for human behaviour into a simple transmission model. Journal of The Royal Society Interface 21:20240038.

Sattenspiel, L., and K. Dietz. 1995. A structured epidemic model incorporating geographic mobility among regions. Mathematical Biosciences 128:71–91.

Saucedo, O., and J. H. Tien. 2022. Host movement, transmission hot spots, and vector-borne disease dynamics on spatial networks. Infectious disease modelling 7:742–760.

Schnyder, S. K., J. J. Molina, R. Yamamoto, and M. S. Turner. 2025. Understanding Nash epidemics. Proceedings of the National Academy of Sciences 122:e2409362122.

Song, H., C. H. C. Hsu, B. Pan, and Y. Liu. 2024. How COVID-19 has changed tourists’ behaviour. Nature Human Behaviour 9:43–52.

Stechemesser, A., M. Kotz, M. Auffhammer, and L. Wenz. 2023. Prolonged exposure weakens risk perception and behavioral mobility response: Empirical evidence from Covid-19. Transportation Research Interdisciplinary Perspectives 22:100906.

Sternberg, E. D., and M. B. Thomas. 2014. Local adaptation to temperature and the implications for vector-borne diseases. Trends in Parasitology 30:115–122.

Stockmaier, S., N. Stroeymeyt, E. C. Shattuck, D. M. Hawley, L. A. Meyers, and D. I. Bolnick. 2021. Infectious diseases and social distancing in nature. Science 371:eabc8881.

Stoddard, S. T., B. M. Forshey, A. C. Morrison, V. A. Paz-Soldan, G. M. Vazquez-Prokopec, H. Astete, R. C. Reiner Jr, S. Vilcarromero, J. P. Elder, E. S. Halsey, et al. 2013. House-to-house human movement drives dengue virus transmission. Proceedings of the National Academy of Sciences 110:994–999.

Stoddard, S. T., A. C. Morrison, G. M. Vazquez-Prokopec, V. Paz Soldan, T. J. Kochel, U. Kitron, J. P. Elder, and T. W. Scott. 2009. The role of human movement in the transmission of vector-borne pathogens. PLoS neglected tropical diseases 3:e481.

Tellier, R. 2022. Covid-19: The case for aerosol transmission. Interface Focus 12:20210072.

Van Der Linden, S. 2022. Misinformation: Susceptibility, spread, and interventions to immunize the public. Nature Medicine 28:460–467.

Ventura, P. C., A. Aleta, F. A. Rodrigues, and Y. Moreno. 2022. Modeling the effects of social distancing on the large-scale spreading of diseases. Epidemics 38:100544.

Verelst, F., L. Willem, and P. Beutels. 2016. Behavioural change models for infectious disease transmission: a systematic review (2010–2015). Journal of The Royal Society Interface 13:20160820.

von Seidlein, L., G. Alabaster, J. Deen, and J. Knudsen. 2021. Crowding has consequences: Prevention and management of covid-19 in informal urban settlements. Building and environment 188:107472.

Wang, W., Y. Nie, W. Li, T. Lin, M.-S. Shang, S. Su, Y. Tang, Y.-C. Zhang, and G.-Q. Sun. 2024. Epidemic spreading on higher-order networks. Physics Reports 1056:1–70.

Wang, Z., M. A. Andrews, Z.-X. Wu, L. Wang, and C. T. Bauch. 2015. Coupled disease–behavior dynamics on complex networks: A review. Physics of Life Reviews 15:1–29.

Wardle, J., S. Bhatia, M. U. Kraemer, P. Nouvellet, and A. Cori. 2023. Gaps in mobility data and implications for modelling epidemic spread: a scoping review and simulation study. Epidemics 42:100666.

Weinstein, S. B., J. C. Buck, and H. S. Young. 2018. A landscape of disgust. Science 359:1213–1214.

Weitz, J. S., S. W. Park, C. Eksin, and J. Dushoff. 2020. Awareness-driven behavior changes can shift the shape of epidemics away from peaks and toward plateaus, shoulders, and oscillations. Proceedings of the National Academy of Sciences 117:32764–32771.

Wesolowski, A., N. Eagle, A. J. Tatem, D. L. Smith, A. M. Noor, R. W. Snow, and C. O. Buckee. 2012. Quantifying the impact of human mobility on malaria. Science 338:267–270.

Weston, D., K. Hauck, and R. Aml ôt. 2018. Infection prevention behaviour and infectious disease modelling: a review of the literature and recommendations for the future. BMC Public Health 18:336.

White, L. A., J. D. Forester, and M. E. Craft. 2018. Disease outbreak thresholds emerge from interactions between movement behavior, landscape structure, and epidemiology. Proceedings of the National Academy of Sciences 115:7374–7379.

Wilber, M. Q., A. Yang, R. Boughton, K. R. Manlove, R. S. Miller, K. M. Pepin, and G. Wittemyer. 2022. A model for leveraging animal movement to understand spatio-temporal disease dynamics. Ecology Letters 25:1290–1304.

Yabe, T., B. G. B. Bueno, X. Dong, A. Pentland, and E. Moro. 2023. Behavioral changes during the COVID-19 pandemic decreased income diversity of urban encounters. Nature Communications 14:2310.

Zhang, H.-F., Z. Yang, Z.-X. Wu, B.-H. Wang, and T. Zhou. 2013. Braess’s paradox in epidemic game: Better condition results in less payoff. Scientific Reports 3:3292.

Zhang, M., S. Wang, T. Hu, X. Fu, X. Wang, Y. Hu, B. Halloran, Z. Li, Y. Cui, H. Liu, Z. Liu, and S. Bao. 2022. Human mobility and COVID-19 transmission: A systematic review and future directions. Annals of GIS 28:501–514.

Zhang, X., F. Scarabel, K. Murty, and J. Wu. 2023. Renewal equations for delayed population behaviour adaptation coupled with disease transmission dynamics: A mechanism for multiple waves of emerging infections. Mathematical Biosciences 365:109068.

Zheng, J. 2020. SARS-CoV-2: An emerging coronavirus that causes a global threat. International Journal of Biological Sciences 16:1678–1685.

Zhou, J., G. Xiao, S. A. Cheong, X. Fu, L. Wong, S. Ma, and T. H. Cheng. 2012. Epidemic reemergence in adaptive complex networks. Physical Review E 85:036107.

